# Heterogeneous epigenetic regulatory patterns link mammalian aging, development, and mortality

**DOI:** 10.64898/2026.09.08.750031

**Authors:** Stanislav Tikhonov, Sergey E. Dmitriev

## Abstract

Aging is often described as a monotonic accumulation of cellular damage, yet all-cause mortality follows a U-shaped trajectory with age, suggesting non-monotonic molecular changes. We investigated links between early childhood development, aging, and chronic diseases by analyzing DNA methylation in mammalian blood. A meta-analysis of 16 human chronic diseases revealed heterogeneous methylation signatures that formed 2 major disease clusters distinguished by their associations with development and sex-related methylation changes. Although epigenetic entropy increased monotonically across the lifespan, several diseases reduced blood DNA methylation entropy independently of blood cell composition. Across mammals, many CpG sites, particularly in intergenic regions, followed U-shaped age-related methylation changes that paralleled mortality curves. Based on these patterns, we developed epigenetic clocks that predict expected mortality across species and tissues and are effective in detecting a range of disease models. Overall, our findings reveal fundamental links between epigenetic regulation during development, aging, and chronic diseases.

## INTRODUCTION

The aggravation of the primary hallmarks of aging, such as genomic instability and telomere attrition, is currently viewed as progressive and cumulative, suggesting a monotonic deleterious effect of the accumulated cellular damage throughout lifespan ^1,2^. According to the ground zero model ^3^, biological aging of the organism starts during mid-embryogenesis and continues until death, which is reflected in age predictions from several epigenetic and transcriptomic clocks ^4,5^.

Yet, the definition of biological age remains debated ^6^. One way to measure it is via all-cause mortality, which follows a U-shaped curve, declining from birth to around 8-10 years in humans and monotonically rising in later life ^7,8^. The leading causes of death for children aged 0-5 years include accidents, congenital malformations, homicide, and infectious diseases, but also malignant neoplasms and cardiovascular diseases ^9,10^. The latter two constitute the leading causes of adult deaths ^11,12^ and, in this context, are viewed as age-related diseases ^13^, i.e., as manifestations of aging. Progeroid disorders should be separately mentioned: while characterized by an accelerated aging phenotype, some of them (e.g. Hutchinson-Gilford progeroid syndrome) may cause death in early to late childhood ^14^.

In contrast to the damage theory of aging ^15^, some theories propose that the late-life persistence of processes beneficial in early life – such as compensatory responses to damage or genetic programs of developmental growth and nutrient sensing – can become life-limiting ^16–18^. The activity of certain developmental genes, such as *IGF1*, is known to increase in childhood and decline in later life ^2^, suggesting the existence of molecular mechanisms capable of regulating U-shaped behavior of certain cellular processes. While non-linear age-related molecular changes have been reported across multiple omics data types in humans ^19,20^, non-monotonic changes mirroring mortality curves have not been studied at a systems level. Overall, the contribution of childhood developmental processes to aging and age-related disease, as well as whether major systemic changes – such as increasing epigenetic entropy – follow monotonic trajectories across the lifespan, remains unclear.

To date, numerous interventions – including diets, small-molecule drugs and gene therapies – were shown to extend lifespan in model organisms ^21–23^. Systems-level analyses of associated molecular patterns, examined alongside male–female and interspecies gene expression differences, revealed a feminizing effect for many classes of interventions ^24^ and both shared and distinct features with cross-species longevity signatures ^25^. In contrast, the molecular basis of chronic diseases is much less clear. Sex hormone therapy has demonstrated efficacy across multiple age-related conditions ^26–28^, and differences in disease incidence between short- and long-lived mammalian species suggest that underlying mechanisms are linked to maximum lifespan ^29^. However, the systems-level associations of sex differences and maximum lifespan signatures with chronic diseases have not been studied using omics data in a meta-analysis spanning diverse chronic conditions.

Assessing the effect of lifespan-extending interventions using survivorship curves can be costly and time-consuming. Epigenetic clocks offer a more efficient alternative for screening such interventions. Recent advances include models that estimate biological age from proxy measures, such as blood plasma concentrations of aging biomarkers, but these clocks are limited to specific species ^30–32^. A recently developed epigenetic clock predicted chronological age across many mammalian species ^33^. However, no existing epigenetic clock predicts expected mortality as a cross-species metric of biological age.

To fill these gaps, we conducted a meta-analysis of 16 human chronic diseases and compared their blood epigenetic signatures to those of aging, childhood development, sex differences, and maximum lifespan of mammalian species. We analyzed associations of DNA methylation (DNAm) levels with mortality and chronological age, characterized CpG sites with U-shaped age-dependent methylation patterns, and developed epigenetic clocks predicting expected mortality as a metric of biological age across various mammalian species and tissues.

## RESULTS

### Human age-related diseases form two distinct clusters of DNAm signatures

To assemble a representative set of human chronic diseases, we aggregated blood DNAm data from 12 case-control studies covering 16 age-related diseases and progeroid syndromes (Fig. 1a). The dataset included conditions affecting diverse tissues, including neurodegenerative and inflammatory bowel diseases. We identified disease signatures by quantifying disease-associated methylation changes in individual CpGs after adjusting for sex and chronological age (Table S1). To examine links between chronic diseases, development, aging, and sex differences, we identified blood DNAm sex-adjusted signatures of early childhood, late childhood, and adulthood aging as well as an age-adjusted sex (male-specific) signature. To analyze chronic disease signatures in evolutionary context, we also identified a signature of maximum lifespan across 35 mammalian species using the blood DNAm data, adjusted for normalized age (i.e. chronological age divided by maximum lifespan) and sex (Fig. 1a).

**Figure 1.**
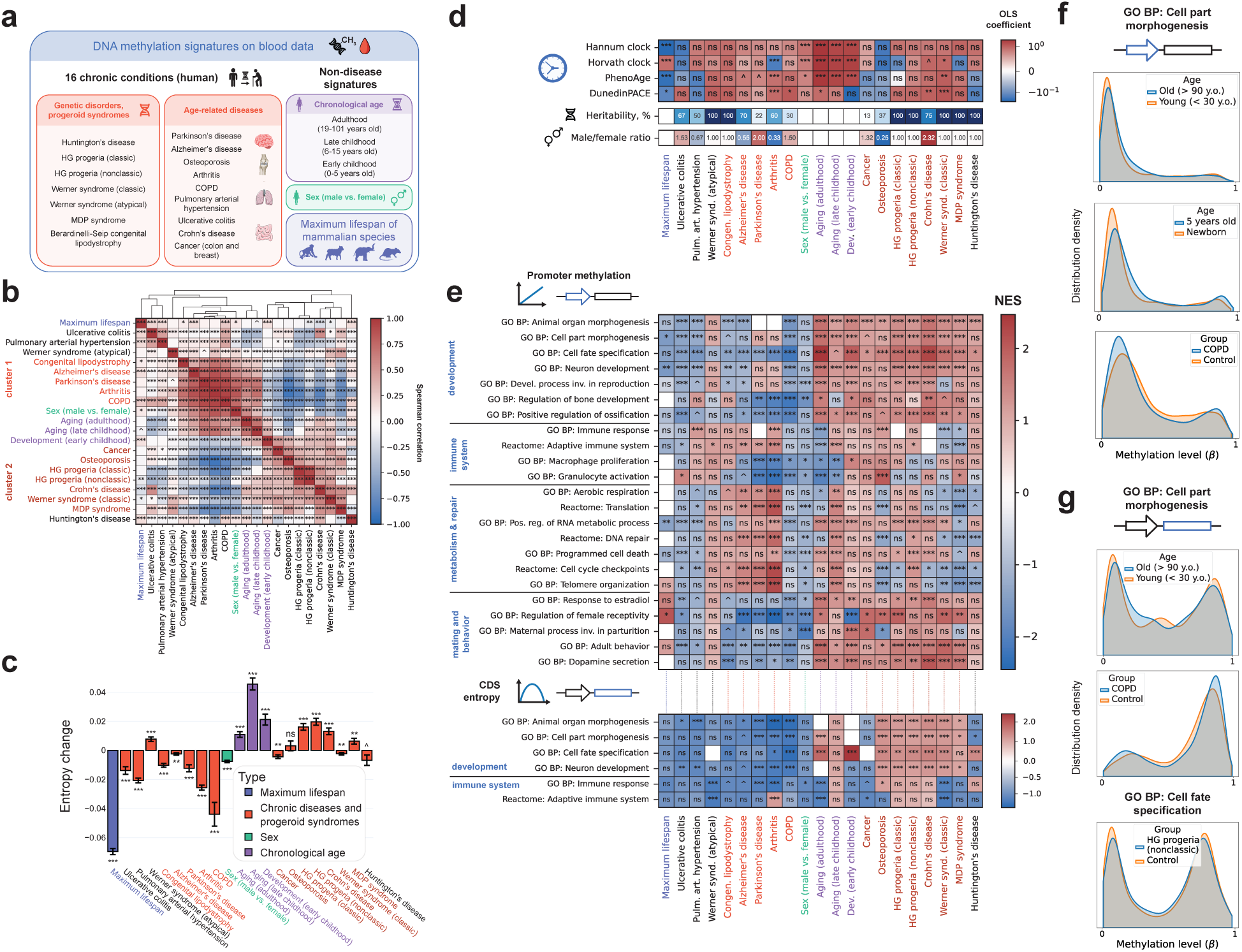
Heterogeneity of DNA methylation profiles of human chronic diseases. **a**, Graphical description of the chronic disorder dataset and the list of non-disease signatures. Color represents the different phenotypes with respect to which changes in DNAm were analyzed in each signature: presence of samples in the disease group and not in the control group (red), chronological age (purple), sex (whether the sample is from a male subject, green) and maximum lifespan (dark blue). **b**, Spearman correlation of disease and non-disease signatures with hierarchical clustering. Disease clusters highlighted in the plot differ in the sign of their correlation with the development in early childhood and sex signatures. The phenotype and disease cluster color schemes, as well as the order of signatures, are preserved in subfigures c, d and e. **c**, Difference in total DNAm entropy with respect to each disease or other phenotypes. **d**, Changes of epigenetic clock predictions with respect to each disease or other phenotype (positive coefficient represents an increase in predicted age in disease patients compared to controls, or males compared to females, or with increasing chronological age or maximum lifespan), and two disease characteristics: heritability and male-to-female incidence ratio. **e**, GSEA enrichment of functional groups of genes in each signature. Top plot represents enrichment results in signatures capturing linear changes in promoter DNAm; bottom plot represents enrichment results in signatures capturing linear changes in CDS DNAm entropy in a CpG-specific manner. For stylistic reasons, the NES scale of the top plot was truncated at around -2.5, the only NES less than the truncated threshold was that of the “GO BP: Regulation of female receptivity” term enrichment in the signature of arthritis. **f**, Examples of promoter DNAm changes in top 100,000 significant CpG sites according to regular DNAm change signatures. **g**, Examples of CDS DNAm changes in top 100,000 significant CPG sites according to DNAm entropy change signatures. ns – adjusted p-value ≥ 0.1 (not significant); ^ – adjusted p-value < 0.1, * – adjusted p-value < 0.05; ** – adjusted p-value < 0.01; *** – adjusted p-value < 0.001.

Overall, the signatures of chronic diseases appeared to be heterogeneous. Spearman correlation analysis revealed that while most (10/16) disease signatures had a positive correlation with the adulthood chronological age signature, correlations with developmental DNAm changes and sex-difference revealed two distinct disease clusters (Fig. 1b, Table S2). These two clusters were robust to both bootstrap resampling of the CpG sites in each signature (Fig. S1a) and to changes in the number of top CpG sites ranked by p-value that were used for the calculation of correlations (Fig. S1b), suggesting that they are not methodological artifacts. For reader’s convenience, we will henceforth refer to cluster 1 as MND (Masculinizing, Negative correlation with Development) and to cluster 2 as FPD (Feminizing, Positive correlation with Development). Notably, the correlation signs of disease signatures with the sex-difference signature largely persisted when restricting the analysis to males or females (Fig. S2), with no significant difference in correlations between male and female disease signatures (Wilcoxon p-value 0.28).

### Epigenetic entropy changes induced by chronic diseases

A well-established hallmark of aging ^2^, the age-dependent increase in DNAm entropy, was confirmed in our analysis for all ranges of chronological age after birth in human blood, including development (Fig. 1c). Notably, not all chronic disorders shared this hallmark, e.g. patients with Huntington’s disease (HD) had lower DNAm entropy compared to age-matched healthy participants (Table 1). Disease-associated entropy changes largely agreed with the identified clusters: most FPD diseases increased entropy, while most MND diseases decreased it. Moreover, males also exhibited significantly lower baseline DNAm entropy than females, consistent with the masculinizing profile of MND diseases. The above-stated results were robust when restricting analyses to promoter or coding DNA sequence (CDS) regions (Fig. S3-S4).

**Table 1.** Summary of entropy and methylation changes for disease clusters, aging, development, and sex signatures. ↓ indicates a (majority) decrease, ↑ indicates a (majority) increase, and “-“ indicates that the sign of change is mixed or unclear due to limited statistical power.

|  | Total entropy | Promoter methylation |  | CDS entropy |  |
| --- | --- | --- | --- | --- | --- |
|  |  | Developmental genes | Immune genes | Developmental genes | Immune genes |
| Cluster 1 (MND) diseases | ↓ | ↓ | - | ↓ | - |
| Cluster 2 (FPD) diseases | ↑ | ↑ | - | ↑ | - |
| Aging | ↑ | ↑ | ↓ | ↑ | ↓ |
| Development | ↑ | ↑ | ↑ | ↑ | - |
| Sex (male vs. female) | ↓ | ↓ | ↓ | - | ↓ |

To validate a significant decrease in blood DNAm entropy for HD, we expanded this analysis to HD models in mice and sheep. Animals’ signatures of HD were positively correlated with the human signature, suggesting overall comparability of the HD-associated DNAm changes across species (Table S3). DNAm entropy significantly decreased in mouse blood in HD model samples compared to controls, but increased in two areas of mouse brain (cortex and striatum, see Fig. S5a,b), suggesting that the effect on entropy may be organ-specific, and that the entropy decrease in blood may be a compensatory response to the HD-induced cellular damage in the brain. Notably, an HIV infection – an example of a condition whose primary affected tissue is the blood – resulted in an increased blood entropy (Fig. S5c).

Diseases in both clusters appear to be diverse. Cluster 2 (FPD) included more genetic disorders and showed higher mean heritability than cluster 1 (MND) (Fig. 1D), although the difference was not significant (Mann-Whitney U test p-value=0.31). Despite opposite correlations with the sex signature, clusters showed no difference in male-to-female incidence ratios (Fig. 1d, Mann-Whitney U test p-value=1). Potential confounding factors, such as chip platform, blood fraction, or the lack of adjustment for sex, could not explain the differences between the clusters (Table S1). Furthermore, predicted blood cell type proportions (based on cell type deconvolution, see Materials and Methods) were largely similar across studies (Fig. S6a), and neither mean predicted cell type proportions, nor the disease-associated shifts in them could account for cluster separation (Fig. S6a,b). Spearman correlations between changes in predicted cell type proportions (Fig. S6b) and entropy (Fig. 1c) were non-significant for all cell types (p.adjusted>0.2), indicating that disease-, age-, and sex-associated entropy changes were not driven by blood cell composition.

Notably, not all chronic diseases had a positive correlation with aging in adulthood – osteoporosis, cancer, pulmonary arterial hypertension (PAH), and ulcerative colitis are examples of diseases with significant negative correlations with the aging signature (Fig. 1b), which further supports the existence of compensatory responses in the blood to age-related damage triggered by diseases that affect other tissues. In addition to this, 6 diseases (e.g., PAH and ulcerative colitis) correlated positively with maximum lifespan of species (Fig. 1b), and all of such diseases resulted in lower DNAm entropy in blood (Fig. 1c), suggesting this particular compensatory response to be a potential natural selection target in long-lived species.

Most established epigenetic clocks detected significant age acceleration during development and aging and in males versus females, but often failed to capture disease-associated acceleration. Although many diseases showed a non-significant trend toward increased DNAm age relative to controls, some (e.g., osteoporosis) showed the opposite trend (Fig. 1d), which is also reflected in low ROC AUC values (see “Pan-mammalian epigenetic clocks of expected mortality”). These results indicate that current epigenetic clocks have limited sensitivity for chronic disease prediction.

To analyze functional changes within disease clusters, we performed Gene Set Enrichment Analysis (GSEA) on both regular DNAm signatures and CpG-wise entropy signatures (see “Materials and Methods”). Promoter regions showed more significant normalized enrichment scores (NES) for DNAm signatures, whereas entropy signatures yielded more significant NES in CDS regions, suggesting more directional, regulated methylation changes in promoters and more stochastic changes in CDS regions (Fig. S7; χ^2^ test of independence for the promoter/CDS × regular DNAm/entropy signature contingency table, p-value 0.0024). Mean promoter DNAm levels negatively correlated with mean expression of the corresponding genes in healthy adult human blood (Spearman ρ=-0.13, p-value 2.9·10^-42^), indicating that promoter methylation enrichment may reflect transcriptional regulation.

### Disease clusters differ in methylation of developmental genes

GSEA revealed a clear distinction between the clusters in methylation change patterns in developmental genes (Table 1). Developmental gene sets showed the largest number of significant NES for both promoter DNAm and CDS entropy signatures (Fig. 1e, top plot). Promoter methylation of developmental genes decreased in response to MND diseases and in males compared to females, but increased with aging, development, and in response to FPD diseases (Fig. 1e, top plot, and Fig. 1f). CDS entropy of developmental genes similarly distinguished the clusters: MND diseases decreased entropy, while FPD diseases increased it (Fig. 1e, bottom plot, and Fig. 1g), consistent with our previous results (Fig. 1c; Fig. S4b). Notably, while DNAm entropy generally increased with age in the majority of CpGs and in most large functional groups of genes during adulthood, we found that methylation entropy of immune genes was in fact decreasing with age (Fig. S7b,d), presumably highlighting a manifestation of inflammaging, an additional regulated compensatory response to age-related cellular damage. Accordingly, the expression levels of the developmental, immune, and other gene sets from Fig. 1e were high in adult human blood (Supplementary Note 1).

Interestingly, we found no major epigenetic pattern that would be common for all 16 chronic diseases. The largest number of disease signatures in which some functional groups of genes were enriched significantly (p.adjusted<0.05) and with the same sign was 7 out of 16 diseases (43.8%). The distribution of the number of CpG associations with diseases was very close to binomial, suggesting that the associations were largely independent across diseases (Fig. S8a,b). Although the meta-signature of all 16 diseases did have 91,688/274,191 significant CpG sites after the Benjamini-Hochberg (BH) correction, it did not represent epigenetic changes common to all diseases but rather the average or most frequent ones: the average disease signature correlation decreased by 0.006 after filtering CpGs by significance in the meta-signature (Wilcoxon test p-value=0.54), and the meta-signature itself correlated negatively with many diseases (Fig. S9).

To elucidate the correlations reported in Fig. 1b, we created two cluster-specific disease meta-signatures. Unlike the global meta-signature, these did capture general trends in DNAm changes within each cluster (Fig. S10). We then identified gene sets with concordant and discordant DNAm changes between the disease meta-signatures and non-disease signatures, which revealed contrasting correlations of disease clusters with development being driven by developmental genes, whereas the co-directional DNAm changes of both clusters with aging arising from different pathways: immune genes for cluster 1 and developmental genes for cluster 2 (Table 2, Fig. S11). The contrasting correlations with the sex signature were also driven by different gene sets (Table 2, Fig. S11).

**Table 2.**
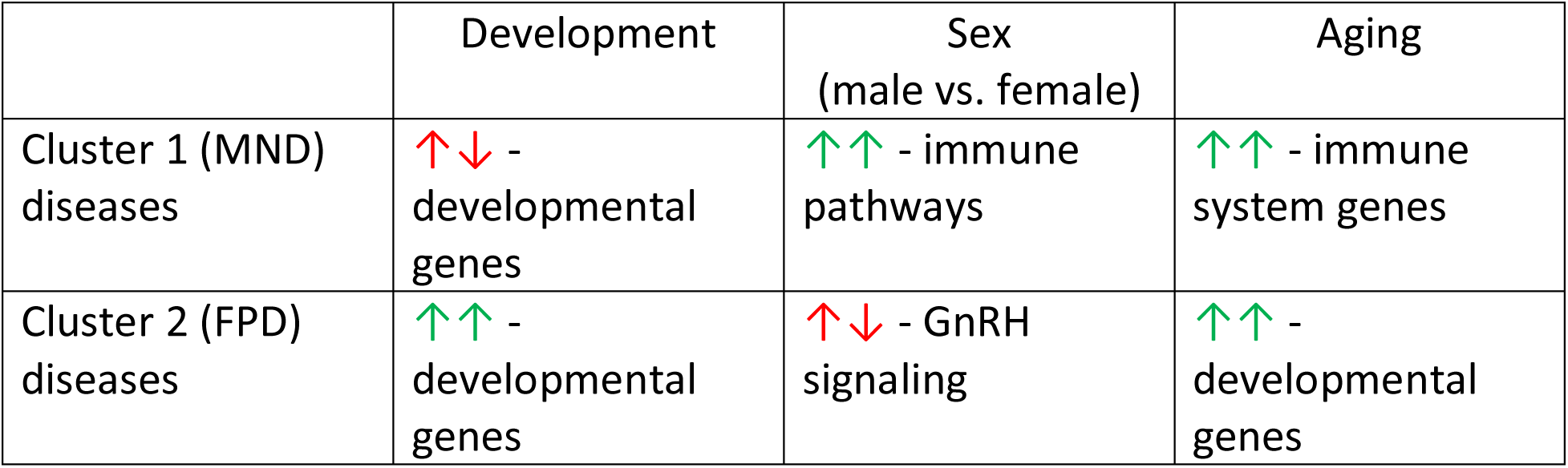
Summary of the groups of genes behind the positive and negative associations of disease clusters and non-disease human signatures according to the results of Fisher’s exact tests. ↑↓ stands for negative correlation and opposite-sign methylation changes, ↑↑ stands for positive correlation and same-sign methylation changes.

Overall, our meta-analysis revealed high heterogeneity of DNAm profiles of chronic diseases, demonstrating mixed-sign associations of disease signatures with development and male *vs*. female differences. We also discovered a number of compensatory responses to age-related cellular damage that were not previously described: decreasing blood DNAm entropy in response to multiple chronic diseases, as well as decreasing DNAm entropy in immune genes with age in adulthood.

### Entropy is shaped by age, sex, and species lifespan

Analysis of associations of non-disease signatures revealed a significant decrease of baseline DNAm entropy in blood of long-lived species after correcting for normalized age and sex (Fig. 1c). It complements previous data showing that the rate of entropy gain per unit of time is inversely associated with maximum lifespan in mammals ^34^. Together with the increase of entropy with chronological age, this suggests a general association between epigenetic entropy and mortality, both across ages and species. To examine it further, we restricted analysis to species showing significant positive correlations between mean DNAm entropy and age (17 species, examples in Fig. S12a).

This analysis confirmed the inverse relationship between baseline DNAm entropy and maximum lifespan (Fig. S12e, Supplementary Note 2) and confirmed a hyperbolic relationship between the rate of change of mean DNAm entropy with age (in years) and maximum lifespan (Fig. S12b–c), in agreement with a previous study by Horvath et al. ^35^. Additionally, when age was divided by each species’ maximum lifespan, the rate of DNAm entropy increase was no longer significantly associated with lifespan (Fig. S12f and S12g), suggesting that mammalian species accumulate a similar overall amount of DNAm entropy over their lifespan, regardless of how long they live. Lastly, we confirmed lower baseline entropy in males compared to females on the filtered dataset (p-value 6.3·10^-5^), and found no male-to-female difference in the rates of entropy change with age and with maximum lifespan (Supplementary Note 3).

### U-shaped trajectories of mammalian DNA methylation and mortality

To further examine links between development, aging, and mortality, we analyzed all-cause mortality, which changes in opposite directions during development and adult aging ^7,8^. We collected survivorship data for 42 species across 11 mammalian orders spanning a wide range of lifespans (Fig. 2a), with sex-specific data for many species. Siler mortality models were fitted for each species and sex pair (Fig. 2b) and validated by strong correlations between predicted and observed maximum lifespan (Fig. S13a). Resulting hazard curves noticeably varied across species and sexes, with many showing elevated childhood mortality (Fig. 2c).

**Figure 2.**
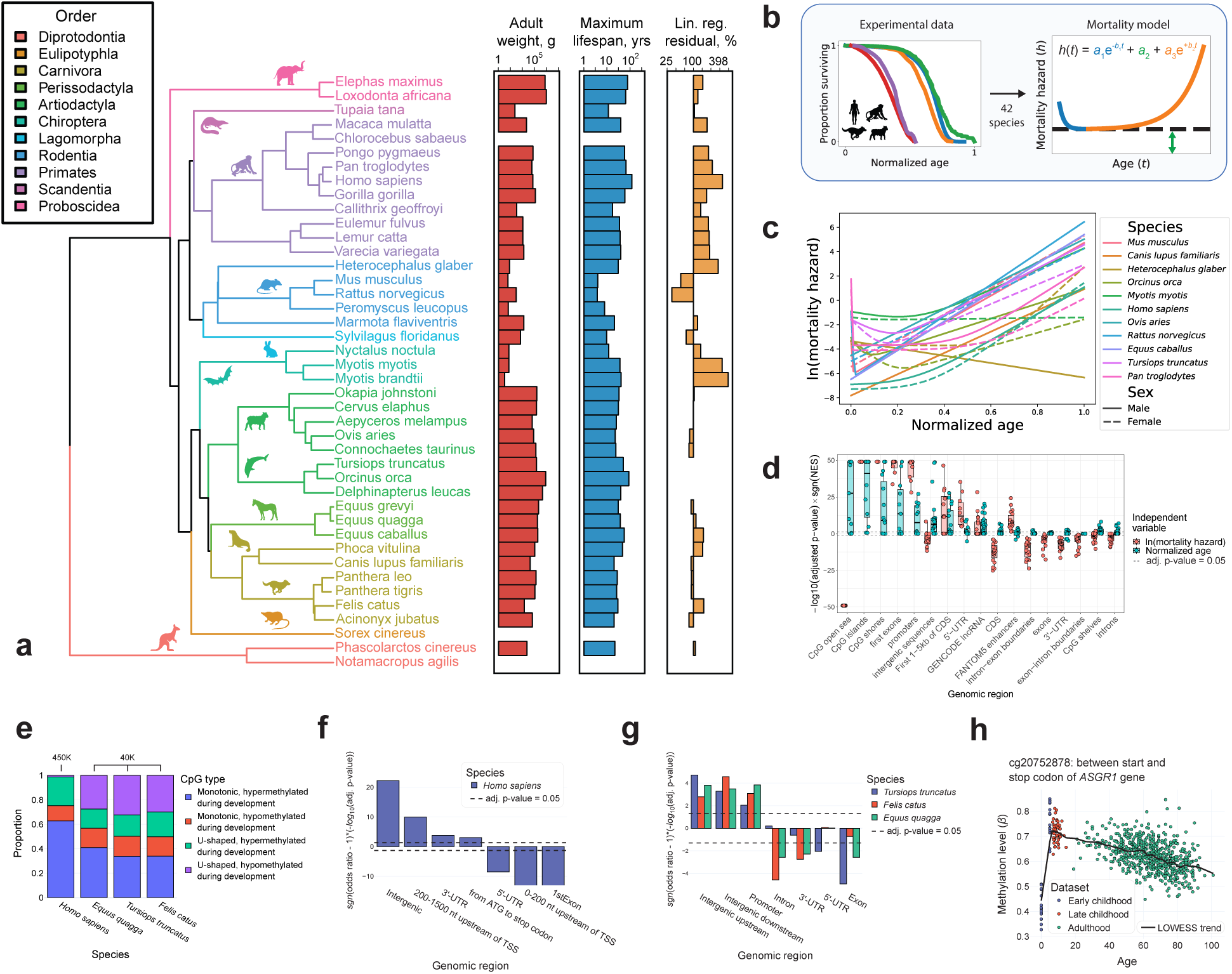
DNA methylation associations with expected mortality across species. **a**, Phylogenetic tree of all species, for which we aggregated mortality data and DNAm data. Side plots show average adult weight (log scale), maximum lifespan (log scale) and the residual from maximum lifespan vs. average adult weight linear regression from AnAge database. **b**, Graphical description of mortality model fitting. **c**, Different shapes of species- and sex-specific mortality hazard curves across mammalian species. **d**, GSEA results for genomic region enrichment in CpG sites whose methylation was associated with normalized age and mortality in various tissues. Dots represent combinations of tissue and sex for which the enrichment scores could be successfully calculated. **e**, Proportions of CpG sites whose methylation levels’ variation with age exhibits monotonic and U-shaped behavior in blood data from 4 mammalian species. A distinction is made between chip platforms: human results came from a different platform compared to the other 3 species. **f**, Fisher tests for intersections of genomic regions with CpG sites whose methylation levels’ variation with age exhibits U-shaped behavior in humans. **g**, Fisher tests for intersections of genomic regions with CpG sites whose methylation levels’ variation with age exhibits U-shaped behavior in *Felis catus*, *Equus quagga* and *Tursiops truncatus*. **h**, An example of a CpG site whose methylation level variation with age exhibits U-shaped behavior in humans.

DNAm data used for this analysis spanned 29 tissues, with most data being from whole blood (Fig. S13b), and covered the whole normalized age scale (Fig. S13c) ^36^. GSEA enrichment of genomic regions in data from multiple tissues revealed that not all regions are equally associated with aging and mortality (Fig. 2d; Fig. S13d). While DNAm status of promoters had a positive association with both normalized age and expected mortality, other regions, such as CpG open sea, intergenic regions, and CDS, were also hypermethylated with age but hypomethylated in individuals with higher mortality rate. Some CDS regions (e.g., exons) showed mortality-associated hypomethylation without a significant association with chronological age in most tissue-sex combinations. Notably, these discrepancies largely disappeared after omitting childhood mortality terms from the mortality models (Fig. S13e), suggesting that some age-related methylation changes counteract the developmental changes (Table 3).

**Table 3.** Proportions of CpG sites with U-shaped age-related methylation changes with respect to the age of minimum mortality among the CpG sites whose methylation level changes significantly (adjusted p-value < 0.05) both before and after that age. Confidence intervals (CI) are Wilson score intervals for binomial proportions.

|  | Proportion of CpG sites with U-shaped methylation changes | 95% CI (lower) | 95% CI (upper) |
| --- | --- | --- | --- |
| *Homo sapiens* | 24.8% | 24.4% | 25.2% |
| *Equus quagga* | 43.2% | 42.5% | 44.0% |
| *Tursiops truncatus* | 49.8% | 49.1% | 50.6% |
| *Felis catus* | 50.2% | 49.5% | 50.9% |

In order to further investigate contrasting DNAm dynamics in development and aging, we identified CpG sites whose methylation in blood significantly changed in opposite directions during these periods in *Homo sapiens*, *Felis catus*, *Equus quagga*, and *Tursiops truncatus* (species with sufficient infant DNAm data). In all four species, there were substantial proportions of CpGs with U-shaped age-related methylation trajectories (Fig. 2e, Table 3, Supplementary Note 4), which significantly overlapped between the 3 non-human species (all pairwise Fisher test adjusted p-values < 10^-292^) and were enriched for intergenic regions in all 4 species (Fig. 2f,g). In humans, they were also associated with CDS regions, genes pertaining to various immune pathways (Fig. S14a), and human endogenous retrovirus (HERV) transcriptional units and active LINEs (long interspersed nuclear elements) (odds ratio 1.6, p.adjusted 0.003). Examples of CpGs exhibiting U-shaped age-related methylation changes are given in Fig. 2h and Fig. S13f. The corresponding genes, *ASGR1* (cg20752878) and *NFIX* (cg17344906), were expressed in blood throughout the whole lifespan (CPM (counts per million) > 1.76 for all samples aged 2-91 years) without significant differential expression in either childhood or adulthood (adjusted p-values > 0.4). *ASGR1* encodes an asialoglycoprotein receptor, which was identified as a therapeutic target in hypercholesterolemia ^37^, whereas *NFIX* encodes a transcription factor essential for myogenesis and hematopoiesis, that was also linked to various types of cancer ^38^. Similar methylation trajectories with significant U-shaped changes were observed in two independent datasets for both CpG sites.

U-shaped patterns were also observed in predicted monocyte and CD4+ T cell proportions in human blood (Fig. S6b), suggesting partial contribution from cell composition. Correlation analysis showed that aging in adulthood and late childhood has an overall masculinizing effect on blood DNAm, whereas early childhood development has a feminizing effect (Fig. 1b). This is consistent with age-related mortality changes, since males generally tend to have higher mortality compared to females ^39^. These correlations may partly reflect blood cell-type changes, particularly the changes of the monocyte and CD4+ T cell proportions (Fig. S6b, Table 4). We observed the predicted monocyte proportion increasing with age in adults, decreasing during development and being greater in adult males compared to females, while the predicted CD4+ T cell proportion displayed the opposite associations (in agreement with experimental results ^40–43^). The observed masculinizing/feminizing effects of aging and development also agree with the corresponding promoter methylation changes in immune genes (Fig. 1e, Table 4). While the positive correlation of sex differences with adult aging can indeed be attributed to similar DNAm changes in immune genes, the negative correlation with development arises mainly from opposite changes in developmental genes (Fig. S11g,i), indicating distinct genomic contributions.

**Table 4.** Summary of feminizing changes in development and masculinizing changes in aging. ↓ indicates a (majority) decrease, ↑ indicates a (majority) increase.

|  | Blood monocyte proportion | Blood CD4+ T cell proportion | Promoter methylation in immune genes |
| --- | --- | --- | --- |
| Aging | ↑ | ↓ | ↓ |
| Development | ↓ | ↑ | ↑ |
| Sex (male vs. female) | ↑ | ↓ | ↓ |

As in human blood, DNAm entropy increased during infant development in all of the three non-human species, with a statistically insignificant increase in *Tursiops truncatus* (Fig. S13g). Overall, this indicates that the entropy is likely to increase in all mammalian species from birth until death.

In summary, we showed that a significant portion of CpG sites demonstrate U-shaped age-related methylation changes in multiple mammalian species, which is confirmed by our analysis of DNAm associations with mortality.

### Pan-mammalian epigenetic clocks of expected mortality

The presence of U-shaped age-related trajectories in both DNAm and mortality indicate that epigenetic clocks of expected all-cause mortality can be developed. Using the mortality and DNAm data described earlier, we constructed and validated epigenetic clocks that predict all-cause mortality as a metric of biological age across various mammalian species (Fig. 3a), including blood-specific and multi-tissue models. The target variable, *ln(mortality hazard)*, had a nearly bell-shaped distribution in both cases (Fig. S15a). We compared elastic net (EN) linear regression, SVM, random forest, and gradient boosting models; nested cross-validation identified SVM as best-performing for both blood-specific and multi-tissue clocks (Fig. S15b,c). Given the interpretability and strong performance of the linear model (mean test-set R² > 0.8), we retained it alongside SVM for further analyses. Examples of out-of-sample performance on healthy human blood are presented in Fig. 3b,c.

**Figure 3.**
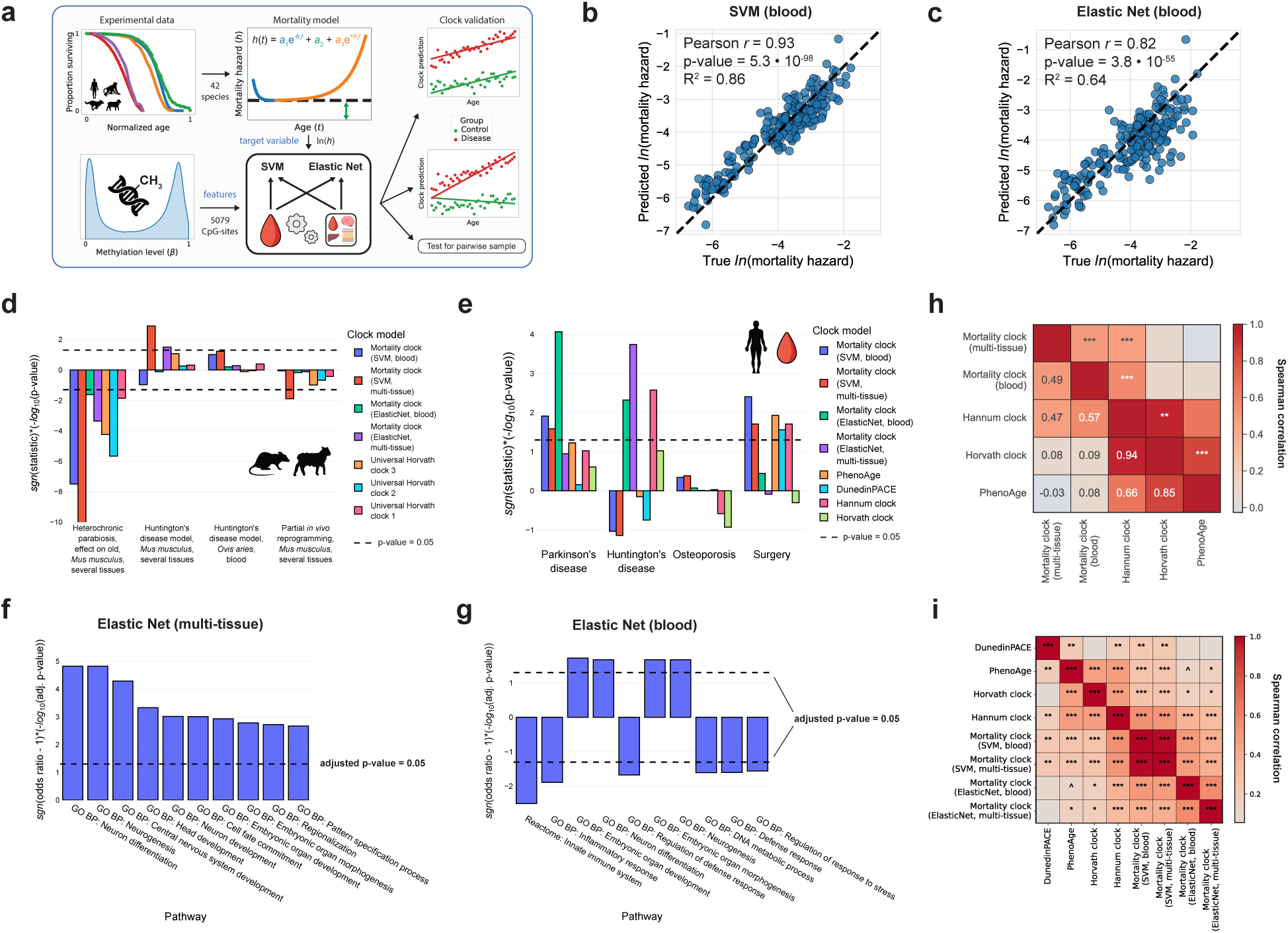
Development and application of epigenetic clocks that predict expected mortality as a metric of biological age across mammals. **a**, Data acquisition, clock training and validation flowchart. **b**, True vs. predicted mortality hazard on out-of-sample healthy human blood for blood SVM, logarithmic scale. True mortality hazard was obtained from chronological age using a sex-specific Siler model. **c**, True vs. predicted mortality hazard on out-of-sample healthy human blood for blood elastic net linear model, logarithmic scale. True mortality hazard was obtained from chronological age using a sex-specific Siler model. **d**, Clock validation on various tissues of *Mus musculus* and *Ovis aries*. “Statistic” represents the effect of a condition (the corresponding OLS coefficient or the median difference in clock predictions in the case of pairwise tests). **e**, Clock validation on human blood. The definition of “statistic” is equivalent to the one used in Fig. 3d. **f**, Fisher test for functional groups of genes’ intersections with top 10% of CpGs according to the multi-tissue elastic net clock’s feature importance. **g**, Fisher test for functional groups of genes’ intersections with top 10% of CpGs according to the blood elastic net clock’s feature importance. **h**, Spearman correlations of linear model coefficients of mortality clocks and benchmark human epigenetic clocks. **i**, Spearman correlations of clock predictions adjusted for age and sex on healthy human blood out-of-sample data. ns – adjusted p-value ≥ 0.1 (not significant); ^ – adjusted p-value < 0.1, * – adjusted p-value < 0.05; ** – adjusted p-value < 0.01; *** – adjusted p-value < 0.001.

We validated the mortality clocks using case-control studies of chronic diseases and other conditions known to affect biological age. We used both data from human blood and data from various tissues of mice and sheep. Clocks were considered effective in detecting a condition if their predictions were significantly higher in the disease group (or equivalent in terms of higher expected biological age) compared to the control group after correcting for age, sex and sometimes other covariates such as tissue or mouse strain (see Table S6). Because some conditions primarily shifted baseline predicted age whereas others altered its rate of increase (Fig. S16a,b), we applied different validation tests tailored to baseline versus rate effects, depending on the condition (Fig. 3a; see Materials and Methods).

Overall, the best models of mortality clocks by mean test-set R^2^ outperformed pan-mammalian chronological clocks on mouse and sheep data (Fig. S17a, Wilcoxon p-adjusted < 0.1 for the multi-tissue SVM vs. all benchmark clocks) and showed comparable performance on human blood (Fig. S17b). Mortality clocks better detected the effects of heterochronic parabiosis (the rejuvenation effect in old mice as well as the pro-aging effect and recovery after detachment in young mice), partial *in vivo* reprogramming, and exercise in mice, as well as HD model in mice and sheep (Fig. 3d, S17c, Tables S7 and S8). Additionally, mortality clocks detected the effect of high fat diet in mice (Fig. S17c).

On human blood data, mortality clocks outperformed human benchmark clocks in predicting Parkinson’s disease (PD), HD, osteoporosis and the effects of surgery and recovery after surgery (Fig. 3e, Fig. S17d, Table S7). Mortality clocks were uniquely capable of detecting PD (Fig. 3e). They were also the only clocks able to detect osteoporosis correctly, although with marginal statistical significance (Fig. 3e and S16b, p-value = 0.06 for the blood SVM mortality clock). Notably, this result could only be obtained when the childhood mortality was present in the training data. The performance gains of mortality clocks with respect to all benchmark clocks were robust to bootstrap resampling of validation datasets (all Wilcoxon p-adjusted < 10^-4^, Table S7). All results of clock comparisons on human, mouse, and sheep datasets are presented in Table S8. The ROC AUC values are also shown on Fig. S18.

The top 10% of CpG sites with the highest feature importances in the multi-tissue and blood EN mortality clocks significantly overlapped with developmental genes (Fig. 3f and 3g, respectively). Linear model coefficients significantly positively correlated with those of the Hannum clock, but not with the Horvath clock or PhenoAge (Fig. 3h). On out-of-sample data from healthy human blood, the age- and sex-adjusted predictions of all clocks used in this study positively correlated with each other (Fig. 3i). As expected, the unadjusted predictions of all clocks also significantly positively correlated with chronological age on out-of-sample data (Spearman correlation p-adjusted < 10^-4^ for all clocks).

Overall, mortality clocks demonstrated broad applicability to both human DNAm data as well as data from model organisms. Their superior performance over pan-mammalian clocks in mice and sheep, together with comparable performance to human-specific clocks on human data, indicates the applicability of mortality clocks for detecting health-modulating interventions on model organisms.

## DISCUSSION

In this study, we showed that chronic diseases have heterogeneous effects on blood DNAm. Global blood DNAm entropy increases monotonically from birth to death in humans, cats, zebras, and likely across all mammals. Nonetheless, a significant proportion of chronic diseases analyzed in this study were associated with decreased blood entropy. This pattern may reflect a compensatory (adaptive) response of the organism to cellular damage in an organ or tissue specifically affected by the disease.

In line with this view, previous studies have shown that not all age-related changes in gene expression and DNAm are harmful ^25,44^. Thus, the expression of some gene sets changes in the same direction with age and in response to lifespan-extending interventions ^25^. It was also suggested that certain age-related compensatory mechanisms may be selected for in long-lived species ^25^. In our study, all diseases whose signatures positively correlated with the maximum lifespan signature were also associated with decreased DNAm entropy in blood, implicating entropy reduction as a potential target of natural selection in long-lived species.

Both decreases and increases of blood DNAm entropy associated with chronic diseases were localized predominantly in developmental genes, both in their promoters and CDS. This was the also the main factor behind the clusterization of DNAm signatures of chronic diseases. Together with significant age-related changes in blood DNAm of developmental genes in adulthood, these results highlight the importance of developmental processes in aging and chronic disease. Their continued regulation outside embryonic development and infancy is consistent with ongoing developmental processes in adulthood, such as neurogenesis and bone remodeling ^45,46^, and with evidence that global developmental regulators such as *Hox* genes remain active beyond development and can regulate behavior, regenerative processes, and neuroplasticity ^47–50^. Blood-mediated regulation of the aforementioned organ-specific processes in adulthood has also been reported (see ^51^ and references therein), but little is known about the role of blood DNAm regulation of developmental genes in chronic disease. Further study is needed to elucidate these associations.

Promoter hypomethylation and decrease of DNAm entropy in developmental genes in blood observed in cluster 1 (MND) diseases may reflect adaptive responses to decreased developmental gene activity in the tissues primarily affected by these diseases. For example, both Alzheimer’s disease (AD) and PD adversely affect adult neurogenesis ^52^, and an impaired neurogenesis or cognitive function has been shown to also be concomitant with all 3 of the remaining cluster 1 diseases ^53–55^. In addition, all cluster 1 (MND) diseases are known to be associated with bone loss ^56–59^ except Berardinelli-Seip congenital lipodystrophy, which increases bone mass ^60^ (congenital lipodystrophy was the most distant member of cluster 1, see Fig. 1b). The importance of these additional symptoms can be highlighted by the effectiveness of the treatment of concurrent bone loss by stimulating bone remodeling against the neurodegenerative symptoms of AD ^57,61,62^. The observed decrease in CDS DNAm entropy and promoter methylation of developmental genes in blood could have a beneficial effect, potentially ameliorating these symptoms. Although the ability of blood immune cells to regulate bone remodeling and adult neurogenesis via cytokine secretion has been reported ^63,64^, the mechanism of such an effect involving epigenetic regulation of developmental genes in blood cells has not been previously described. Our hypothesis could be used to augment the hyperfunction theory of aging ^16^: excessive activity of developmental genes in adulthood might be one of the causes of the aging phenotype, but in some cases, the expression of these genes in blood may play a role in beneficial compensatory responses to chronic diseases.

Another feature discriminating cluster 1 diseases from cluster 2 was correlation with the sex signature. Cluster 1 (MND) diseases had a masculinizing effect on blood DNAm, while cluster 2 (FPD) diseases had a feminizing effect. This finding partly agrees with the therapeutic success of hormonal therapy of the corresponding diseases, but the results from the literature are mixed. For example, estrogen treatment reduces the risk and alleviates the symptoms of both AD and PD in postmenopausal women ^26,27,65^ and may have beneficial effects against both diseases in men ^66,67^ and in male mice ^68,69^. But testosterone treatment also displayed some beneficial effects against AD symptoms in men ^70^, while PD was linked to both high ^71^ and low ^72^ testosterone in men in different studies. The symptoms of osteoporosis and Crohn’s disease can improve both in women treated with estrogens ^73,74^ and in men treated with testosterone ^28,75^. Estrogen deficiency has been linked to an increased risk of rheumatoid arthritis ^76^ in women, but so was testosterone deficiency in men ^77^. The correlations of disease signatures with the sex signature are particularly interesting in the context of previous findings indicating that many of the lifespan-extending interventions have a feminizing effect on mouse gene expression profiles (e.g. caloric restriction and growth hormone deficiency), while certain interventions have a masculinizing effect (e.g. rapamycin) ^24^. While the feminizing or masculinizing effects were sex-specific for some lifespan-extending interventions ^24^, the feminizing or masculinizing effects of chronic diseases were largely similar across sexes.

The connection between the feminizing/masculinizing effects of chronic diseases and their association with development remains unclear. Our results show that early childhood development has a feminizing effect on DNAm in human blood, while aging in adulthood has a masculinizing effect. This pattern can be viewed as a global U-shaped change that parallels mortality dynamics: mortality decreases during early childhood development and increases in adulthood, while males have higher mortality risks compared to females ^39^. Consistent with these findings, blood cell type deconvolution revealed a significant increase of CD4+ T-cell proportions and a decrease in monocyte proportions during early childhood, with opposite changes during adulthood and in males compared to females. Because the adaptive immune function strengthens during early childhood ^78^ and declines with aging ^79^, these patterns may relate to the stronger adaptive and innate immune responses generally observed in adult females compared to males, with higher proportion of CD4+ T cells ^80^. However, changes in blood cell proportions associated with chronic diseases were mostly not significant. Further investigation is needed to elucidate the reasons behind these associations.

The finding of U-shaped behavior in age-related DNAm changes was expected due to many developmental processes having a peak activity at a certain age from early to late childhood, which can be illustrated by age-related changes in the serum concentration of IGF1 ^81^. The age-related increase in blood monocyte percentage that we observed was consistent with previous studies ^40^ and is linked to the development of age-related inflammatory phenotype ^82^. The significant decrease of monocyte proportion during postnatal development could be explained by the active development of adaptive immune system and an increasing proportions of T cells and B cells. The ability of mortality clocks trained on data with childhood mortality to detect primary osteoporosis, which could not be detected correctly by the existing clocks, highlights the importance of U-shaped epigenetic age-related changes.

Several limitations of our study should be noted. First, all human disease and non-disease signatures in our meta-analysis were based on the blood data. Inclusion of data from other tissues, especially the ones corresponding to the origin of pathological processes for each chronic disease, would elucidate the differences between local and systemic deleterious and compensatory changes. We were able to analyze gene expression patterns indirectly via promoter methylation, but further investigation of transcriptomic signatures is needed to shed more light onto the heterogeneous mechanisms of chronic diseases. Second, since chronic diseases are very diverse, accumulation of new data on chronic pathologies and other age-accelerating conditions and its integration into the meta-analysis would make the conclusions more generalizable. Lastly, the analysis of developmental changes and U-shaped patterns was complicated by the scarcity of DNAm data from the earliest months and years of life of different mammalian species. Further study of the developmental changes and U-shaped patterns in age-related DNAm changes could generalize the findings of our work to a wider range of organisms.

## CONCLUDING REMARKS

Our findings revealed substantial heterogeneity in the effects of chronic diseases on blood DNAm, encompassing changes in DNAm entropy, distinct pro-aging changes in immune or developmental genes, and feminizing or masculinizing DNAm shifts. Together, these results highlight the importance of developmental programs and sex differences in the context of aging, chronic disease and mortality. Non-linear age-related changes mirroring mortality dynamics included changes in blood proportions of CD4+ T cells and monocytes, providing a link between age-related changes and sex differences. We observed decreases in blood DNAm entropy in CDS and promoter regions of developmental genes in response to 9 out of 16 analyzed chronic diseases, suggesting a potential compensatory response to tissue-specific cellular damage that could be selected for in long-lived species. Blood DNAm analysis across species revealed decreased baseline blood DNAm entropy in long-lived mammalian species compared to short-lived species, while overall entropy accumulation throughout lifespan was similar across species. Lastly, we developed epigenetic clocks predicting all-cause mortality, whose applicability across species could expand the scope and generalizability of research in the field of aging and lifespan-extending interventions. Overall, our results link intra- and inter-species mortality patterns with heterogeneous epigenetic regulation of aging and chronic disease.

## ACKNOWLEDGEMENTS

We are deeply grateful to Dr. Alexander Tyshkovskiy and Alec Eames for their invaluable contributions and insightful discussions during the initial stages of the study.

## AUTHOR CONTRIBUTIONS

S.T. conceived the study, performed data analysis; S.T. and S.E.D. interpreted the data; S.E.D. supervised the study; S.T. wrote the manuscript with contributions from S.E.D. All authors read and approved the final version.

## MATERIALS AND METHODS

### Meta-analysis of human chronic diseases in the context of aging, early childhood development, sex differences and species longevity

For the meta-analysis of chronic disease, the human blood DNAm data was taken from 12 case-control studies from NCBI GEO ^83,84^: GSE32148 (ulcerative colitis and Crohn’s disease) ^85^, GSE84003 (PAH) ^86^, GSE131752 (classic and atypical Werner syndrome, MDP syndrome) ^87^, GSE214297 (congenital lipodystrophy) ^88^, GSE144858 (AD) ^89^, GSE72774 (PD) ^90,91^, GSE42861 (rheumatoid arthritis) ^92^, GSE118468 (chronic obstructive pulmonary disease (COPD)) ^93^, GSE89093 (cancer, mostly of the colon and breast) ^94^, GSE99624 (primary osteoporosis) ^95^, GSE182991 (classic and nonclassic Hutchinson-Gilford progeria) ^96^ and GSE146917 (HD) ^97^. In the case of HD, the comparison was made between manifest and pre-manifest patients. In the case of PAH, the comparison was made between patients with rheumatoid heart disease with reversible heart valve dysfunction and secondary PAH and controls. For non-disease signature calculation, we used data from the following studies: GSE40279 (sex and adult aging signatures) ^98^, GSE104812 (late childhood aging) ^99^, GSE83334 (early childhood development) ^100^ and GSE223748 (maximum lifespan of mammalian species) ^33,36,101^. In the case of maximum lifespan, we only used DNAm data from blood for the disease meta-analysis. Prior to statistical analysis of maximum lifespan, we removed outlier samples using PCA for species with larger sample sizes and filtered out CpG sites whose methylation level was between 0.45 and 0.55 in more than 10% of samples. Transgenic animal data and embryonic data were filtered out before the analysis. Maximum lifespan data was taken from the AnAge database ^102^, and it was increased in the cases of species for which we had samples with larger chronological ages than the corresponding maximum lifespans reported in AnAge. In the case of naked mole rat, the maximum lifespan was set to 40 years ^103^. The total number of species was 35 (the same species that we used for blood epigenetic clocks). When restricting the correlation analysis to males or females, only the datasets with sex annotation and sufficient samples for each sex were used.

The disease methylation signatures were calculated as vectors of OLS coefficients for disease for all CpG sites in the following regression model:

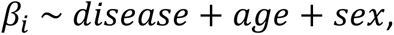

where *β_i_* is the methylation level of i-th CpG, *disease* was equal to 1 for the disease group and 0 for the control group, and *sex* was added to the model where gender data was available (1 corresponds to male and 0 corresponds to female). The entropy signatures were calculated in a similar way with a modified model:

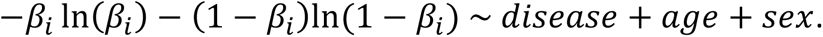

The total (mean) DNAm entropy was calculated as 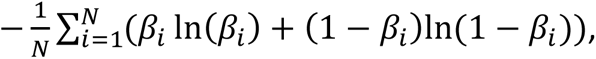 where *N* is the number of CpG sites (the word “total” in the name refers to the fact that the entropy is calculated across many CpG sites).

For total DNAm entropy signatures, we only looked at the top 10,000 CpG sites ranked by p-value in the corresponding regular signatures. In the case of HD models on mouse and sheep, we looked at the top 5,000 CpG sites.

With non-disease signatures, the disease term was not present in the regression models. Instead, the coefficients of min-max-normalized variables of interest were analyzed. In the case of maximum lifespan, the model was

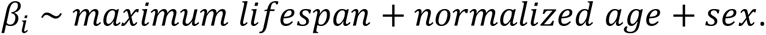

For longitudinal (early childhood development) and twin (cancer) data, the signatures were constructed with one-sample t-tests for the distributions of methylation level differences between later and earlier time points and case and control samples, respectively.

The procedure for the calculation of the signatures of blood cell type proportion changes for Fig. S6b was identical to the one described above, but the methylation levels were replaced with blood cell type proportions (see Materials and Methods, section “Cell-type deconvolution”).

Hierarchical clustering of disease and non-disease signatures was performed with scipy ^104^ using average linkage and (1-*cor*)/2 as distances, where *cor* denotes Spearman correlation calculated using the intersection of top 100,000 CpG sites ranked by p-value of the association of interest (disease/control in disease signatures, age in the signatures of aging and development, sex in the sex signature and maximum lifespan in the maximum lifespan signature). Using the threshold of 100,000 CpG sites ensured large intersection sizes in all pairs of signatures. Disease meta-signatures were calculated by applying the rma.mv function from the metafor ^105^ package in R to the OLS coefficients of the disease term and their standard errors. Prior to this, both the OLS coefficients and their standard errors were scaled by the standard deviation of the coefficients within each disease.

In the consensus clustering analysis, we also used hierarchical clustering with average linkage and (1-*cor*)/2 as distances, where *cor* denotes Spearman correlation calculated using intersections of top *N* CpG sites with the lowest association p-values (*N* depends on the exact procedure, see below). In each clustering, there were 2 cluster labels assigned to disease signatures according to the root node split of the corresponding hierarchical clustering dendrogram. To generate different clusterings, we used two procedures. The first procedure used bootstrap sampling of CpG sites leaving only 30% of sites without replacement and then using *N*=100,000 for correlations. The bootstrap random seed was different across 1000 runs and also for each disease within each run. The second procedure involved iterating over different values of *N* without bootstrap. We used a linear scale from *N*=10,000 to *N*=300,000 with 50 equally spaced values (rounded). With some of the smallest values of *N*, minimum intersections were quite small, potentially leading to more noise in correlations. The minimum, mean, median and maximum intersection sizes for all 50 values of *N* are listed in Table S9. Figure S1 shows consensus index, which is the proportion of clusterings (from 1000 bootstrap clusterings or clusterings with 50 different values of *N*) in which a pair of disease signatures ended up with the same cluster label. We also computed the proportion of clusterings where cluster 1 and cluster 2 (described in Fig.1b) diseases formed their own exclusive clusters at some height cutoff in the hierarchical dendrogram (by exclusive we mean that all of the diseases from cluster 1 (or cluster 2) and no other diseases formed a separate cluster). The 95% confidence intervals (CI) for these proportions were computed as Wilson score intervals for the binomial proportions using proportion_confint function from statsmodels package ^106^ in Python.

The signature of early childhood development was validated using additional data from GSE132181 and GSE103657 ^107^. These studies did not have longitudinal data, so the additional signatures of early childhood development were calculated with linear regression as described above. Coefficients of the intersection of 10,000 most significant CpG sites from the two signatures of postnatal development showed Spearman correlations of 0.84 (p-value 5.2*10^-221^) and 0.93 (p-value 1.8*10^-155^), respectively, suggesting the overall validity of the signature of early childhood development used in the meta-analysis of human chronic diseases.

To estimate the effect size of the divergence between the distribution of the number of disease associations for CpG sites and the theoretical binomial distribution which assumes independent associations, we used the difference in means of the observed and theoretical distributions, Cohen’s ω, a variant of Cramér’s V applicable to the goodness of fit scenario, and פ (Fei) ^108^. Because the number of categories is more than 2 in this case, Cohen’s ω could exceed 1, which is why we added פ (Fei) and the variant of Cramér’s V. The formula for the latter is provided below:

Cramér’s 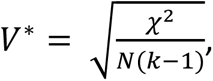, where *N* is the total sample size and *k* is the number of categories (in this case *k* = *n* + 1, where *n* is the number of diseases).

All adjusted p-values in this article were calculated using the Benjamini-Hochberg method ^109^.

To identify gene sets with concordant and discordant DNAm changes for cluster meta-signatures and non-disease signatures, we selected CpGs sites whose methylation changes had the same or opposite signs in a meta-signature and a non-disease signature, filtered them by the respective adjusted p-values, and utilized Fisher tests to find significant intersections of these groups of CpG sites and functional groups of genes.

### Cell-type deconvolution

We estimated blood cell-type proportions from bulk DNAm data using constrained quadratic programming. This approach models bulk methylation (*B*) as a linear combination of cell-type-specific methylation profiles (*X*) weighted by their proportions (*P*): *B* = *X* ∗ *P*. Given the observed bulk methylation (*B*) and reference cell-type methylation profiles (*X*), we solved for cell-type proportions (*P*) using quadratic programming, applying constraints that ensured the proportions were non-negative and summed to one.

We applied platform-specific reference methylation profiles measured on each array type to deconvolute bulk profiles. One reference profile was measured on the Illumina 450K array, and another on the EPIC array. These were used to deconvolute bulk profiles measured on the 450K and EPIC arrays, respectively. This approach accounts for platform-specific technical biases. The reference methylation profiles encompassed six major blood cell types: neutrophils, monocytes, natural killer (NK) cells, B cells, CD4+ T cells and CD8+ T cells. Reference CpG sites were selected based on their differential methylation patterns across cell types, identifying the most distinctively hypermethylated and hypomethylated sites for each cell type.

### CpG site location annotation and analysis

CpG sites’ location (both gene symbols and genomic regions) was taken from chip manifest files for Illumina HumanMethylation450 BeadChip ^110^, Infinium MethylationEPIC ^111^ and Illumina HorvathMammalianMethylChip40 BeadChip ^112^. CpG sites were assigned to promoters if their genomic region was “TSS200” or “TSS1500” or to CDS regions if their region was annotated as “Body” according to the UCSC database. If a CpG-site was not assigned to any gene-related genomic region, it was classified as “Intergenic”. In the case of the maximum lifespan signature, promoter/CDS labeling was done based on the annotation for the human genome hg19.

All gene set enrichment analyses (GSEA) were done with the fgseaMultilevel function from the fgsea ^113^ R package with default parameters. Gene set annotations were taken from GO BP ^114,115^, KEGG ^116–118^, Reactome ^119^ and BioCarta ^120^ databases using msigdbr ^121^ package in R. In order to apply it to CpG sites, we converted the gene sets into CpG sets using chip manifest files for mapping (the choice of the manifest file depended on which chip the DNAm data came from). Fisher tests for intersections of CpG sites and gene sets were done in the same manner using the fisher_exact function from scipy ^104^. For GSEA, the CpG sites were ranked according to their OLS effect coefficient over the effect standard error for disease and other signatures.

A list of CpG sites located in HERV transcriptional units and active LINEs was taken from ^122^.

### Disease heritability and male to female incidence ratios

Heritability data for chronic diseases was taken from the following studies: osteoporosis (bone loss heritability was averaged between spine, hip and radius) ^123,124^, rheumatoid arthritis ^125^, COPD ^126^, ulcerative colitis and Crohn’s disease ^127^, hypertension ^128^, AD ^129^, PD ^130,131^, cancer (average between colorectal and breast cancers) ^132^. Male to female incidence ratios were taken from the following studies: osteoporosis (data from NHANES 2005-2008 study for hip and lumbar spine bone density) ^133^, rheumatoid arthritis (average between the ages below 50 and above 60-70 years old) ^134^, COPD ^135^, ulcerative colitis and Crohn’s disease ^136,137^, hypertension ^138^, AD ^139^, PD ^140^, cancer (colorectal) ^141^.

### Mortality data

All-cause mortality data was taken from the following studies: *Connochaetes taurinus albojubatus* ^142^*, Felis catus* ^143^*, Panthera leo* ^144^*, Tursiops truncatus* ^145^*, Delphinapterus leucas* ^146^*, Notamacropus agilis* ^147^*, Sylvilagus floridanus* ^148^*, Phoca vitulina* ^149^*, Tupaia tana* ^150^*, Marmota flaviventris* ^151^*, Okapia johnstoni* ^152^*, Equus caballus* ^153^*, Callithrix geoffroyi, Eulemur fulvus, Lemur catta, Varecia variegata, Acinonyx jubatus, Panthera tigris, Equus grevyi, Pongo pygmaeus, Macropus fuliginosus* ^154^*, Cervus elaphus* ^155^*, Homo sapiens* ^156^*, Ovis aries* ^157^*, Equus quagga* ^158^*, Pan troglodytes* ^159^*, Orcinus orca* ^160^*, Macaca mulatta* ^161^*, Gorilla gorilla* ^162^*, Loxodonta africana, Elephas maximus* ^163^*, Phascolarctos cinereus* ^164^*, Sorex cinereus* ^165^*, Myotis myotis, Myotis brandtii, Nyctalus noctula* ^166^*, Canis lupus familiaris* ^167^*, Peromyscus leucopus* ^168^*, Chlorocebus sabaeus* ^169^*, Mus musculus* ^170^*, Rattus norvegicus* ^171^*, Heterocephalus glaber* ^103^. In many cases, the mortality data were sex-specific. The data in these studies were most frequently presented as survivorship functions. To obtain mortality hazards, Siler mortality models ^172^ were fitted to the experimental data, allowing for omission of terms representing juvenile or senescence-associated mortality hazards if such models resulted in lower mean squared errors (MSE) compared to the full model. Parameter optimization was performed using the optim function in R. To obtain robust results, initial values for all parameters of the mortality models were set to a grid of 6 values from 10^-3^ to 10^2^ (log scale) for all parameters, after which the model with the lowest MSE was chosen.

### Phylogenetic tree

The phylogenetic tree for 42 mammalian species was constructed using TimeTree ^173^. The average adult weight, maximum lifespan and the residual from the regression of log(maximum lifespan) against log(mean adult weight) were taken from AnAge. The species were assigned to mammalian orders according to the NCBI taxonomy database ^174^.

### DNA methylation associations with mortality and chronological age

For GSEA analysis of DNAm associations with mortality and chronological age, CpG sites were ranked with -sgn(coefficient)*log_10_(p-value) from the linear regression of the methylation level against ln(mortality hazard) or normalized age with a correction for sex. Since the GSEA analysis often resulted in NaN values due to unbalanced ranked lists, tissue- and sex-based subsets of the data from GSE223748 were analyzed from all tissues, blood, brain, skin, muscle, liver, kidney, and both sexes, male and female (we included all combinations of these tissue and sex subsets). For this analysis, we used genomic region annotation from the annotatr ^175^ package for R. Prior to this analysis and all other statistical analysis of GSE223748, we filtered out CpG sites whose methylation level was between 0.45 and 0.55 in more than 10% of samples to ensure a bimodal distribution of methylation levels.

CpG sites with a U-shaped pattern of methylation level change with age were defined as the sites whose methylation levels changed in the opposite directions with adjusted p-values < 0.05 in the development and adulthood aging signatures. For human, these signatures were calculated from GSE83334 and GSE40279, respectively. For *Felis catus*, *Equus quagga* and *Tursiops truncatus* both signatures were calculated using the blood data from GSE223748 (both sexes), after dividing it into development and adulthood using the age of minimum mortality hazard (averaged between males and females). The monotonic CpG sites were identified in the same way but they had the same signs of methylation change in development and adulthood. The methylation trajectories of cg20752878 and cg17344906 were verified using additional childhood data from GSE103657 and healthy adult samples from GSE72774. Both in these age signatures and in the original age signatures based on GSE83334 and GSE40279, the p-values for the above-stated CpG sites were < 0.001.

### Gene expression analysis

For gene expression analysis we used the data from GSE25507 (peripheral blood lymphocytes from human children, microarray) ^176^ and GSE134080 (whole blood from adult humans, RNA-Seq) ^177^. Microarray data was preprocessed as follows. array probe IDs were mapped to Entrez gene IDs, then intensities were averaged within each gene. The resulting intensities were log-transformed and scaled. In the case of GSE25507, disease group samples were filtered out as the first step of the preprocessing. RNA-Seq data was preprocessed in the following way. The GSE134080 data had Ensembl gene IDs, which were mapped to Entrez IDs and the counts were summed within each Entrez ID. Then only the genes that had at least 10 reads in at least one third of samples were kept, the rest were filtered out. The counts were subsequently normalized with the RLE method ^178^ and log-transformed. For differential expression with age calculation, limma ^179^ was used in both cases. Differential expression was calculated with GSE134080 for adulthood and with GSE25507 for development, and in the case of development, only the samples up to 7 years old were used. For calculating the correlation of promoter methylation and gene expression, GSE134080 data was used, where the Entrez IDs were mapped to gene symbols and the normalized and log-transformed expression values were averaged across samples. Only the Entrez IDs with a 1-to-1 mapping to gene symbols were kept in this case. GSEA was then applied to a list of gene symbols from GSE134080 ranked by the mean normalized log-transformed expression.

Additionally, to calculate expression levels throughout lifespan, healthy control samples from GSE279448 (peripheral blood mononuclear cells, RNA-Seq) were used. CPM values for *i*-th gene in *j*-th sample were calculated as: 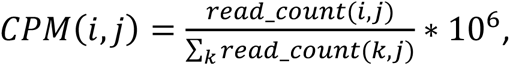 where the denominator is the library size for the *j*-th sample.

### Epigenetic clocks

We used 4 regression models for training the epigenetic clock to predict all-cause mortality: linear model with elastic net regression (ElasticNet function), SVM with nonlinear kernels (SVR function) and random forest (RandomForestRegressor function) from scikit-learn ^180^ and gradient boosting from xgboost ^181^. The clocks were trained to predict mortality in blood and in all tissues, but because the fitting times were longer for all tissues, we only used the linear model (as the most interpretable) and SVM (as the model with the highest R^2^) in this case. The natural logarithm of the mortality hazard was used as the target variable. To facilitate the cross-platform applicability of the resulting clocks, we used the methylation levels of 5079 CpG sites that corresponded to the intersection sites covered by Illumina HumanMethylation450 BeadChip, Infinium MethylationEPIC and Illumina HorvathMammalianMethylChip40 BeadChip as features. The majority of the training data was from GSE223748 (processed in the same way as for maximum lifespan signature calculation but without the CpG site filtering), but we also added human blood data from GSE40279 and chimpanzee blood data from GSE136296 ^182^. The tissues present in GSE223748 included blood, brain, liver, muscle, kidney, skin, lung, heart, adipose tissue, spleen, pancreas, thyroid, lymph node, spinal cord, bone marrow, and various tissues of gastrointestinal tract, male and female reproductive systems. The total sample size for all tissues was 9981 samples from 42 species from 11 mammalian orders. The sample size for blood was 4076 samples from 35 species from 7 mammalian orders.

For hyperparameter optimization and quality assessment we employed a nested cross-validation procedure: the total sample was split into training and validation (70%) and test (30%) sets 20 times with random shuffling and species stratification, and each time the 70% were further subdivided into training and validation sets with the StratifiedKFold function from sci-kit learn with 10 folds, random shuffling and species stratification. For the multi-tissue SVM model, 7 outer splits instead of 20 were performed due to long training times. The training and validation sets were used for Bayesian hyperparameter optimization with optuna ^183^, after which the models were trained on the joined training and validation sets and their quality was assessed on the test set, resulting in test-set R^2^ distributions for each model type. The final models’ (linear regression and SVM, all tissues and blood-only) hyperparameters were optimized on the whole dataset with 100 trials in optuna. Hyperparameter importances and contour plots for R^2^ in the hyperparameter space were made with optuna (Fig. S15d-f).

For validation of mortality clocks, we used additional datasets with case-control studies of conditions known to affect biological age from blood and other tissues. In the case of human, in addition to the datasets used in the meta-analysis of chronic diseases, we used the data from GSE142536 (major surgery, blood) ^184^. In the case of non-human data, we used GSE147002 (HD model, multiple tissues, *Mus musculus*) ^97^, GSE190665 (partial *in vivo* reprogramming, multiple tissues, *Mus musculus*) ^33,101^, GSE224361 (heterochronic parabiosis, multiple tissues, *Mus musculus*) ^185,186^, GSE147003 (HD model, blood, *Ovis aries*) ^97^, GSE267903 (effect of exercise, *Mus musculus*) ^187^, GSE199979 (high fat diet, *Mus musculus*) ^188,189^. For validation on human data, we compared the mortality clocks to some existing human epigenetic clocks that predict chronological age (Hannum clock ^98^ and Horvath clock ^190^) and biological age (PhenoAge ^30^ and DunedinPACE ^31^). For validation on non-human data, we compared the mortality clocks to the only existing epigenetic clocks that are applicable across species: the pan-mammalian chronological clocks ^33^. The high fat diet dataset was the only dataset where only 78-89% of CpG sites from the features of the benchmark clocks were present (for comparison, only 49% of CpG sites from the mortality clock features were present in this dataset). The rest of the datasets had all CpGs from the features of the benchmark clocks, and in the cases where not all of the 5079 mortality clock features were present, mortality clocks were refitted on their training data using only the CpG sites present in the corresponding validation dataset.

Before validating the clocks, we classified each age-accelerating condition into “intercept”, “interaction” and paired sample. The intercept-interaction distinction was made based on whether the condition had more CpG sites with significantly different baseline methylation levels between study groups or with significantly different rate of change of methylation between the groups. We ran regressions with two different models,

*β_i_* ∼ *condition* + *age* + *sex* (the “intercept” model) and *β_i_* ∼ *condition* ∗ *age* + *age* + *sex* (the “interaction” model, *condition* ∗ *age* denotes the interaction term), for each dataset where the sample was not paired (the cancer dataset had paired twin data and the surgery dataset had paired longitudinal data). For each condition that had more significant CpG sites in the “intercept” signature we used the “intercept” test, for conditions with more significant CpG sites in the “interaction” signature we used the “interaction” test, and for conditions with paired samples we used the Wilcoxon test for clock prediction differences between the condition group and the control group (in the case of surgery, the time point “postoperative day 1” corresponded to the “condition” group and the time points before surgery and “postoperative day 4-7” corresponded to the “control” groups). The “intercept” test was a t-test for the coefficient of the disease term in the following regression:

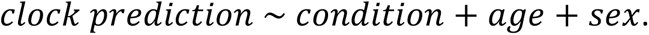

The “interaction” test used a different regression:

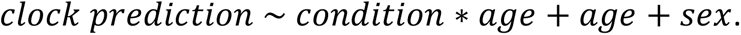

In some datasets, additional batch effects, namely tissue or strain, needed to be accounted for. In such cases, we used a mixed model with a random effect corresponding to the tissue or strain. The resulting test type assignment is listed in Table S6. For tests whose results were presented in Fig. 1D, “intercept” tests were applied to all conditions where the sample was not paired. To standardize the scale of effect sizes for Fig. 1D, we first ran the regression without the term of interest (disease, age, sex or maximum lifespan, depending on the dataset), only with confounding variables, and then ran another regression only with the term of interest and without the confounding variables on the standardized residuals of the first regression. In the cases of paired samples, the clock prediction differences were standardized (but not centered). Prior to running the tests, all methylation levels outside the [0, 1] closed interval were set to NaNs and all NaNs were imputed with mean methylation values for the corresponding CpG sites.

In order to determine which clock was “better” in detecting some condition affecting biological age, we only looked at the statistical significance and the direction of clock prediction change, because a larger magnitude (absolute value) of prediction change does not necessarily indicate a better ability of some clock to differentiate between the case and control groups compared to another clock. We compared signed z-scores (positive z-scores based on p-values of tests multiplied by -1 where a condition’s effect on clocks was in the opposite direction compared to the expected direction) between clocks, computed as 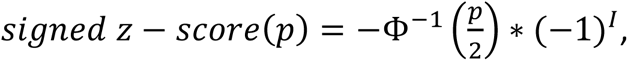, where *p* is the p-value for the condition effect, Φ^−1^(*x*) is the inverse of the cumulative distribution function for the standard normal distribution, and *I* is an indicator that takes the value 1 when the condition’s effect on the predictions of a clock was in the opposite direction compared to the expected direction and the value 0 otherwise. The expected direction was an increase in clock prediction for age-accelerating conditions (e.g. chronic diseases, effect of surgery) compared to the corresponding control groups, and a decrease in clock prediction for age-decelerating conditions (e.g. partial *in vivo* reprogramming, recovery after surgery) compared to the corresponding control groups. In addition to the effect of a condition on clock predictions, the p-value for its coefficient can be affected by the correlation of the clock’s predictions with the covariates (age and sex), but it was generally desirable for the mortality clocks to correlate well with chronological age and differentiate between sexes in addition to differentiating the condition group from the control group, as long as the coefficients for the age and sex (*sex* = 1 for male and *sex* = 0 for female) terms were positive (see Table S8 for the signs of the coefficients corresponding to the age and sex terms).

In order to obtain CIs and a measure of statistical significance for the result of clock comparisons, we employed a bootstrap procedure that preserved at least 3 samples from each of the case and control groups, generating 1000 bootstrap datasets for each condition. For each dataset, the best benchmark clock was chosen based on the observed results prior to bootstrap. For mortality clocks, two options were tested: the clock with the best performance on a given dataset prior to bootstrap and the best model according to the mean test-set R^2^ for the corresponding tissue (blood SVM clock for blood data and multi-tissue SVM clock for data from other tissues or a dataset with several tissues). For each dataset, we computed signed z-scores for the chosen mortality clock and the chosen benchmark clock, computed their difference (mortality clock - benchmark clock) and ran the Wilcoxon test on the bootstrap distribution of such signed z-score differences. The CIs for the signed z-score differences were computed using the same bootstrap distributions. Whenever we say that one clock “outperformed” another clock, we mean that it demonstrated a greater signed z-score and the Wilcoxon adjusted p-value was significant. In the case of human chronic diseases and OLS regression, this would mean that one clock gave a lower p-value for the coefficient of the study group (disease/control) after correcting for age and sex compared to another clock given that both clocks’ study group coefficients were positive. If one clock’s study group were negative in this case, the clock would get a negative signed z-score.

ROC AUC was calculated in the context of patient-control classification or an equivalent classification task in terms of biological age differences (e.g. before and after surgery). Different thresholds for age- and sex-adjusted clock predictions allowed classifying the samples and comparing the result to the true study group labels.

The feature importance was calculated as the absolute values of coefficients in the linear model of mortality clocks. Correlations of mortality clock predictions with chronological age and the predictions of existing human clocks were computed on the DNAm data from the blood of healthy individuals from GSE72774, because GSE40279 was part of the training set for mortality clocks and the Hannum clock.

## SUPPLEMENTARY TABLE LEGENDS

**Table S1.** Overview of the datasets used in the meta-analysis of chronic diseases.

**Table S2.** Description of clusters of chronic conditions.

**Table S3.** Correlations of the signatures of HD models in mice and sheep with the human HD signature, and the effects of the HD models on DNAm entropy.

**Table S4.** GSEA enrichment of functional groups of genes in blood expression.

**Table S5.** GSEA enrichment of the CpG-based leading edges mapped to genes in the mean expression profile of adult human blood. The leading edges were obtained from the GSEA enrichments of the corresponding pathways in the regular DNAm signature of aging (adulthood) and the entropy signature of aging. The signature of aging that the leading edge corresponds to is described in the “Analysis” column.

**Table S6.** Test type classification of the conditions used for epigenetic clock validation.

**Table S7.** Bootstrap results for clock validation.

**Table S8.** Clock validation results.

**Table S9.** Intersection size statistics for different thresholds for the number of top CpG sites ranked by the p-value from a chronic disease or a non-disease signature.

