## Supplementary Notes and Figures for "Heterogeneous epigenetic regulatory patterns link mammalian aging, development, and mortality"

**Supplementary Note 1.** Enriched gene sets from disease GSEA are expressed in human blood.

To validate GSEA findings from Fig. 1e, we checked whether the gene sets with significant NES from Fig. 1e are expressed in adult human blood. We performed GSEA on a mean gene expression profile of adult human blood, revealing significant enrichment of immune, developmental, programmed cell death, translation and other pathways, indicating that the observed DNAm changes occur in genes expressed during adulthood (Table S4).

Additionally, to validate the GSEA associations discovered specifically for the signature of aging in adulthood in Fig. 1e, we looked at the leading edges of immune and developmental gene sets whose promoter DNA methylation levels or CDS entropy were associated with aging in adulthood. In agreement with the previous result, this analysis showed that the leading-edge genes were overall significantly enriched in the mean expression profile from adult human blood and therefore highly expressed compared to the rest of the transcriptome (Table S5).

**Supplementary Note 2.** Baseline DNAm entropy is inversely related to maximum lifespan in mammals.

The intercept coefficients in $entropy \sim age\_in\_years$ regressions did not demonstrate a significant association with maximum lifespan (Fig. S12d), but a similar analysis applied to per-sample DNAm entropy estimates showed that the baseline (age=0) mean DNAm entropy does significantly decrease with increasing maximum lifespan (Fig. S12e), thereby confirming our previous result in Fig. 1c on the filtered dataset (17 species whose entropy significantly positively correlated with age). The regression model used in Fig. S12e fixes the functional relationship between the rate of entropy change per year and maximum lifespan to hyperbolic (because of the normalized age term), and has more statistical power when it comes to determining the functional relationship between the baseline DNAm entropy and maximum lifespan, because it is applied to individual samples, which explains the difference between the results in Fig. S12e and S12d.

**Supplementary Note 3.** Baseline DNAm entropy depends on sex, while the rate of its change with age shows no significant dependence.

The following regression $entropy \sim maximum\_lifespan + normalized\_age + male$, where $male$ is a binary variable that equals to 1 when the sex is male and 0 when it is female, when applied to the filtered dataset (17 species), resulted in a significant negative coefficient -0.004 for the $male$ term (p-value $6.3 \cdot 10^{-5}$), which confirms the dependence of baseline DNAm entropy on sex identified in Fig. 1c. Adding the $normalized\_age * male$ interaction term into the regression above did not result in a significant coefficient (p-value 0.57) and also made the coefficient of the $male$ term less significant (p-value 0.06). We observed the exact same result when adding the same interaction term ($age * male$) into the regression model on human blood data (p-value 0.7 for the interaction term), which means that there is no significant dependence of the rate of entropy change with age (per year or per maximum lifespan) on sex in the data we analyzed. Lastly, adding the $maximum\_lifespan{:}male$ interaction term into the multiple-species regression also resulted in a non-significant coefficient (p-value 0.72), indicating no significant dependence of baseline entropy rate of change with maximum lifespan on sex in the filtered dataset.

**Supplementary Note 4.** CpG sites with U-shaped age-related methylation trajectories.

The proportions of CpGs with U-shaped age-related methylation trajectories was higher than we might expect due to random chance in a sample with generally monotonic behavior: $\chi^2$ goodness-of-fit tests with the expected proportions of 5% and 10% all resulted in p-values < $10^{-307}$. The difference in proportions between humans and the 3 other mammals could be attributed to different chip platforms. While the common genomic region for such CpGs was intergenic in all four species, other genomic regions and functional groups of genes differed in their associations with these CpG sites between the human data and the data from the other 3 mammals (Fig. 2f,g and S14). Notably, positive correlations between all three human chronological age signatures (Fig. 1b) supports the fact that the majority of age-related DNAm changes across the entire lifespan in human blood are monotonic (Table 3).

### SUPPLEMENTARY FIGURES

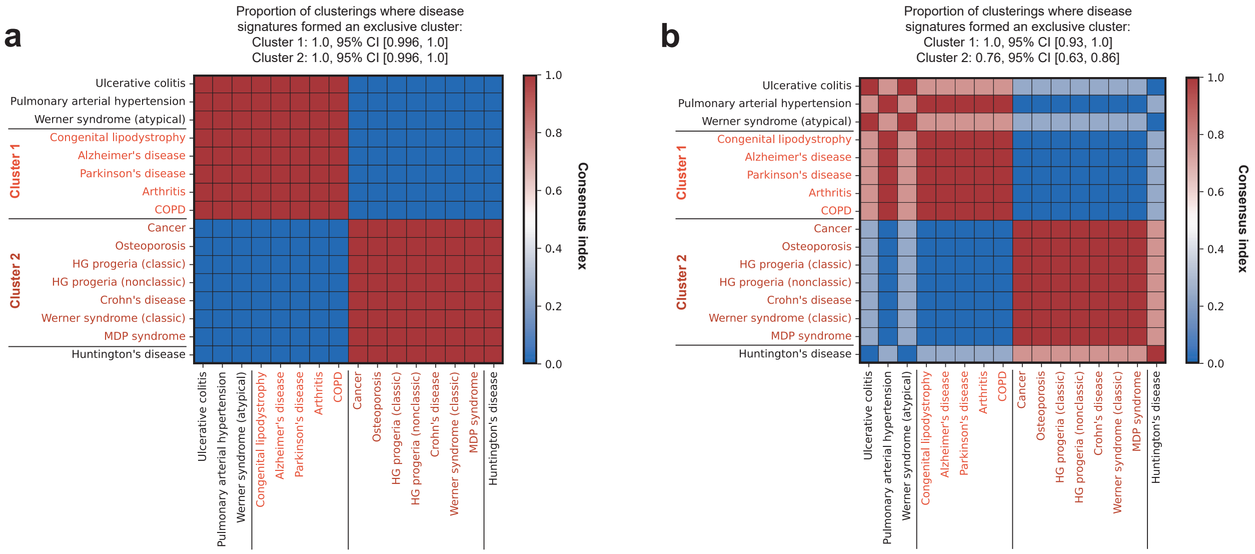


**Figure S1.** Consensus clustering of chronic diseases. Consensus index is defined as the proportion of clusterings in which a pair of disease signatures was assigned to the same cluster. The clustering was hierarchical, and in each run disease signatures were divided into 2 clusters according to the root node split. For details, see Materials and Methods.

**a**, 1000 bootstrap runs with 30% of CpG sites sampled without replacement.

**b**, 50 different thresholds for the number of CpG sites with lowest p-values that are used for correlation calculation (linear scale from top 10,000 sites to top 300,000 sites).

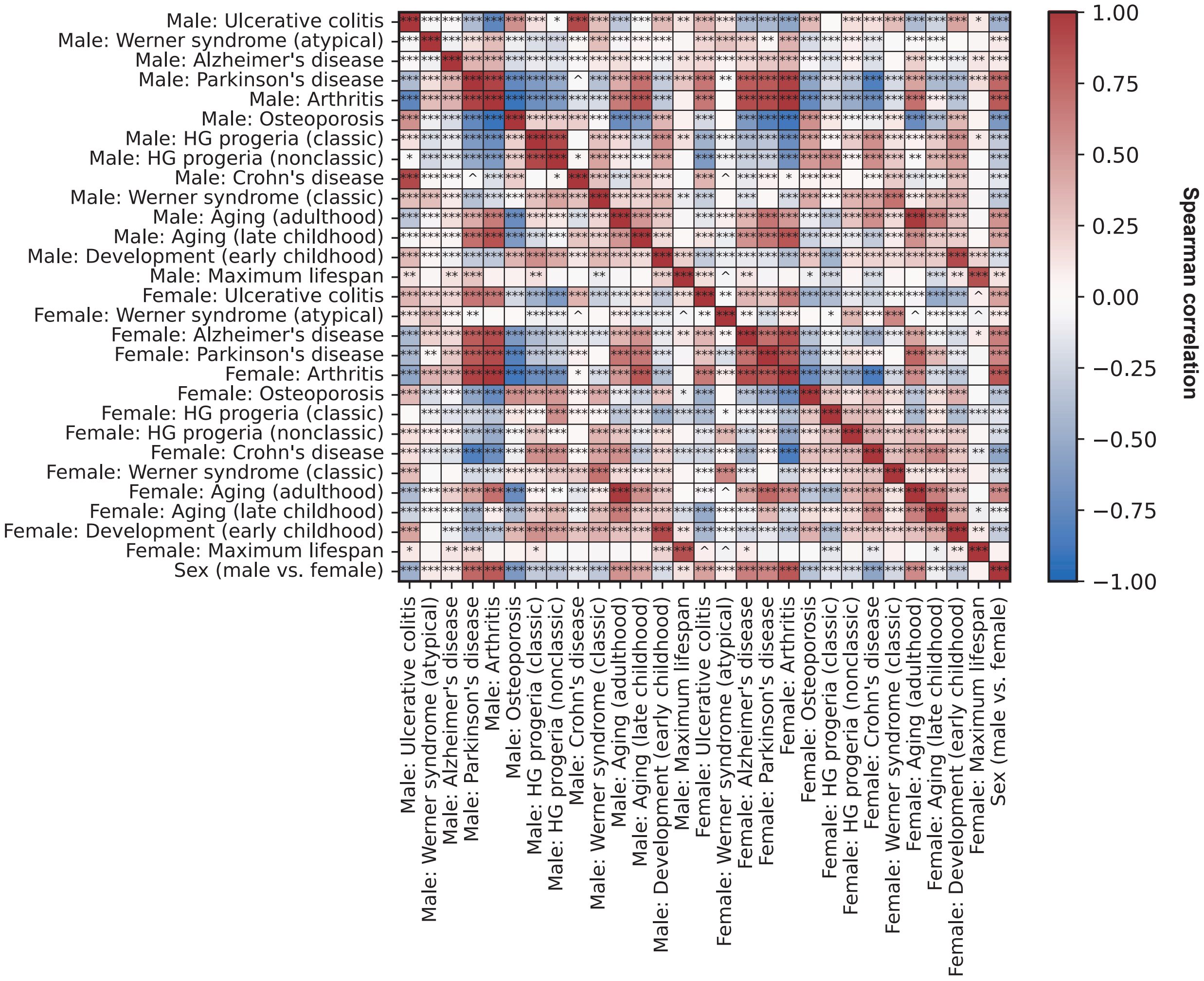


**Figure S2.** Correlations between DNAm signatures of chronic diseases, chronological age, sex differences and maximum lifespan, separately for males and females.

^ – p adjusted < 0.1; * – p adjusted < 0.05; ** – p adjusted < 0.01; *** – p adjusted < 0.001.

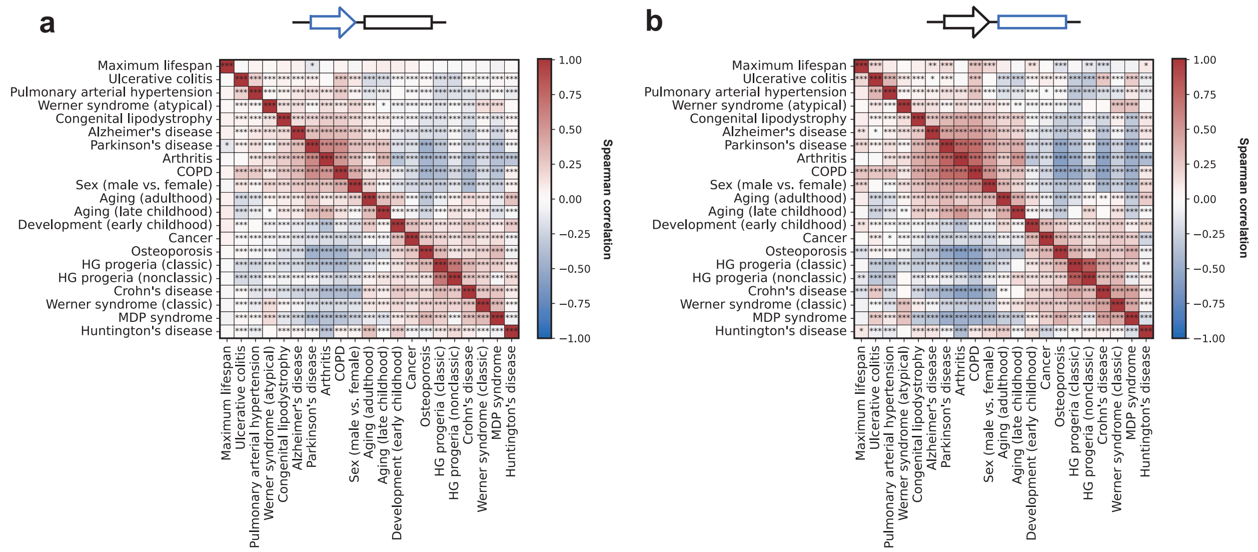


**Figure S3.** Spearman correlations of blood DNA methylation signatures of diseases and non-disease signatures.

**a**, Analysis is restricted to promoter CpG sites.

**b**, Analysis is restricted to CpG sites in CDS regions.

^ – p-value < 0.1, * – p-value < 0.05; ** – p-value < 0.01; *** – p-value < 0.001.

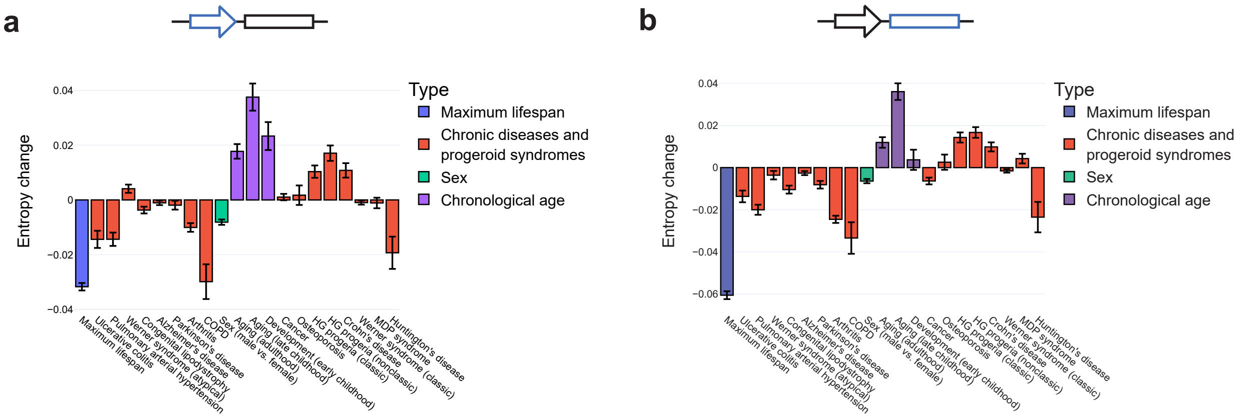


**Figure S4.** Total DNA methylation entropy change in blood with respect to chronic diseases and other phenotypes.

**a**, Analysis is restricted to promoter CpG sites.

**b**, Analysis is restricted to CpG sites in CDS regions.

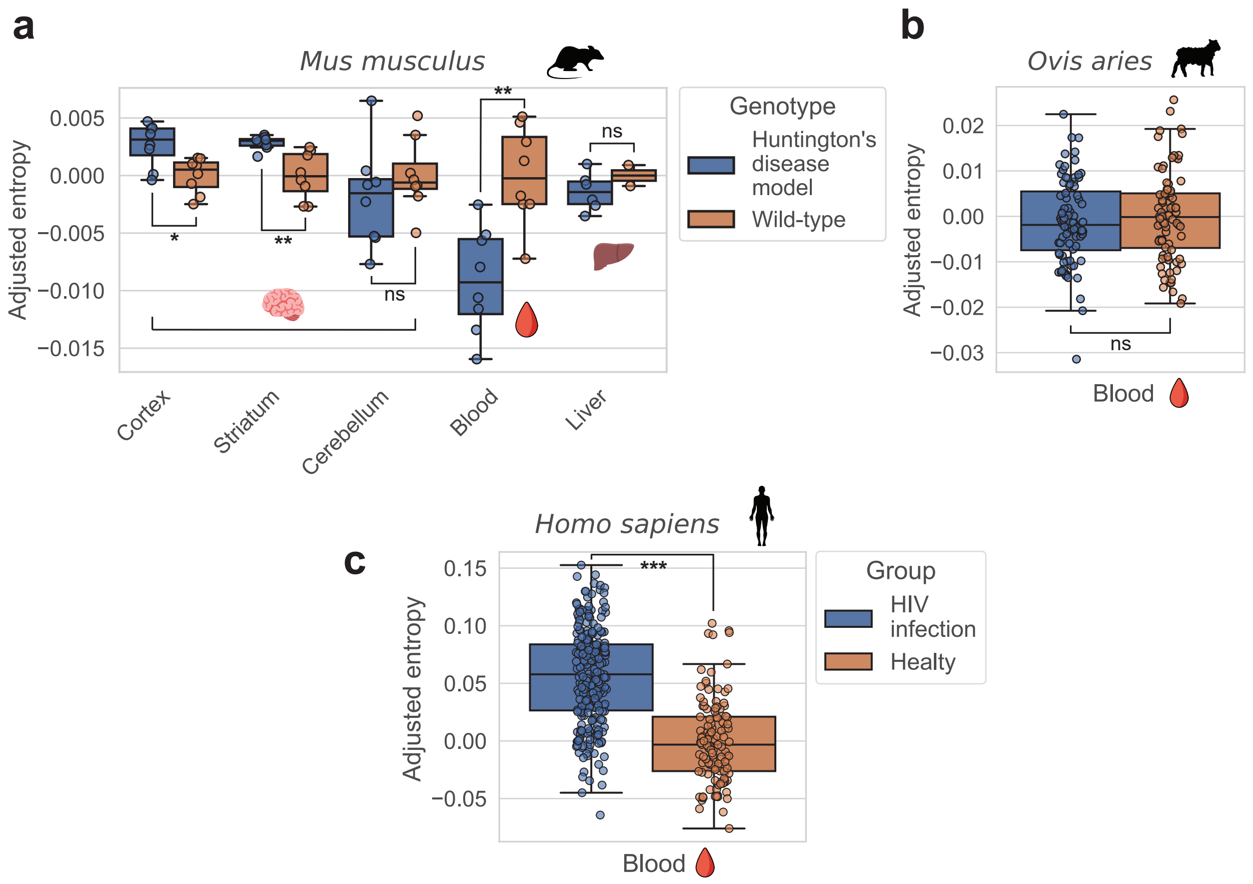


**Figure S5.** Tissue-specific effects of diseases on DNAm entropy adjusted for age, sex and mean per-tissue methylation level.

**a**, Model of Huntington's disease, *Mus musculus*.

**b**, Model of Huntington's disease, *Ovis aries*.

**c**, HIV infection, *Homo sapiens*.

ns – p adjusted ≥ 0.1 (not significant); * – p adjusted < 0.05; ** – p adjusted < 0.01; *** – p adjusted < 0.001.

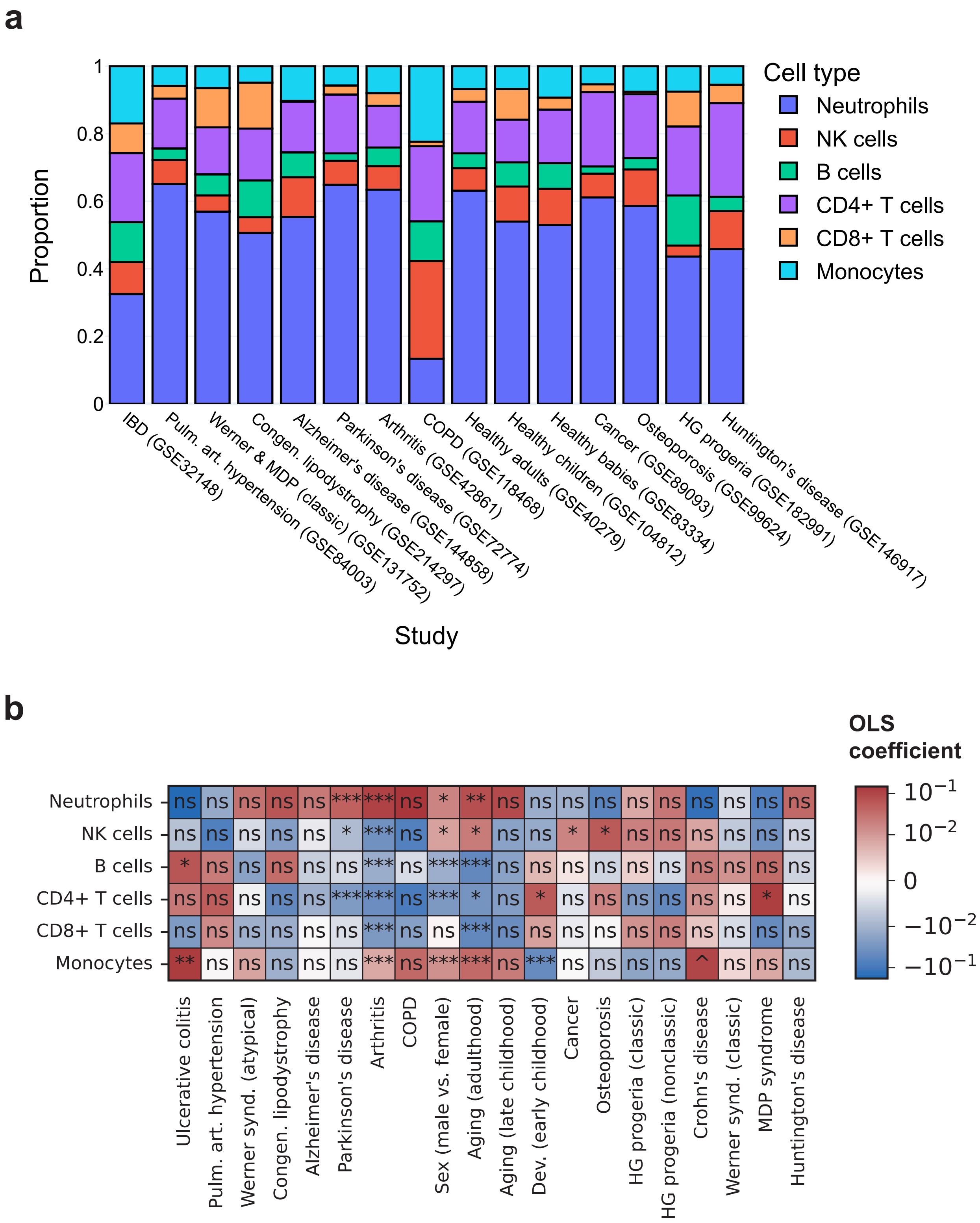


**Figure S6.** Predicted blood cell type proportions and their changes with age and disease.

**a**, Predicted blood cell type distributions for all human studies used in the meta-analysis of human chronic disease.

**b**, Changes in predicted blood cell type proportions with respect to each disease or other phenotype (positive coefficient represents an increase in predicted age in disease patients compared to controls, or males compared to females, or with increasing chronological age).

ns – p adjusted ≥ 0.1 (not significant); ^ – p adjusted < 0.1, * – p adjusted < 0.05; ** – p adjusted < 0.01; *** – p adjusted < 0.001.

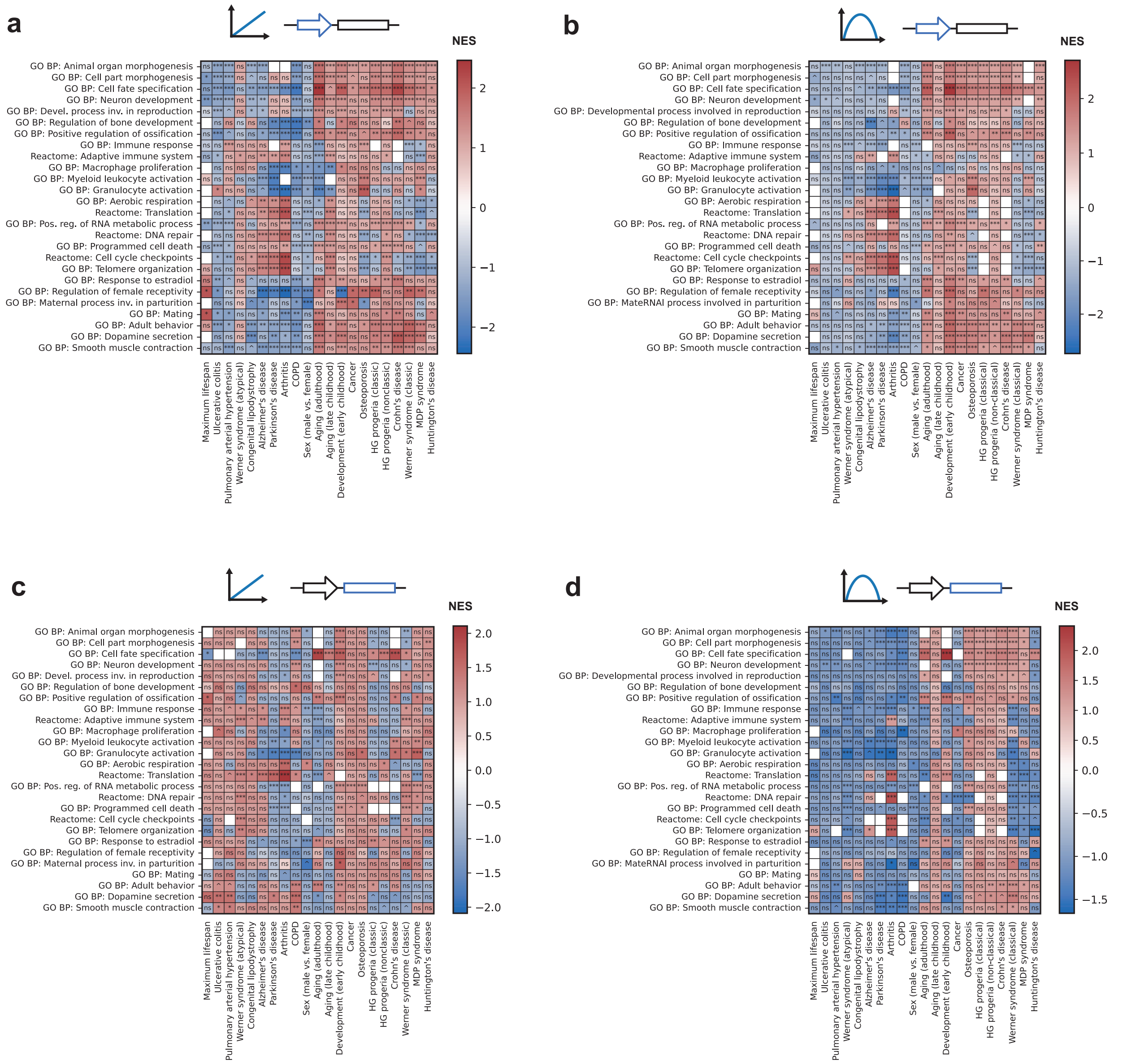


**Figure S7.** GSEA enrichment of functional groups of genes in disease and non-disease signatures of blood DNA methylation and entropy changes in promoters and CDS. The first icon in each subfigure represents the signature type, while the second icon represents the genomic region.

**a**, Signatures are capturing linear changes in promoter DNA methylation.

**b**, Signatures are capturing linear changes in CpG-specific DNA methylation entropy in promoters.

**c**, Signatures are capturing linear changes in CDS DNA methylation.

**d**, Signatures are capturing linear changes in CpG-specific DNA methylation entropy in CDS regions.

ns – p adjusted ≥ 0.1 (not significant); ^ – p adjusted < 0.1, * – p adjusted < 0.05; ** – p adjusted < 0.01; *** – p adjusted < 0.001.

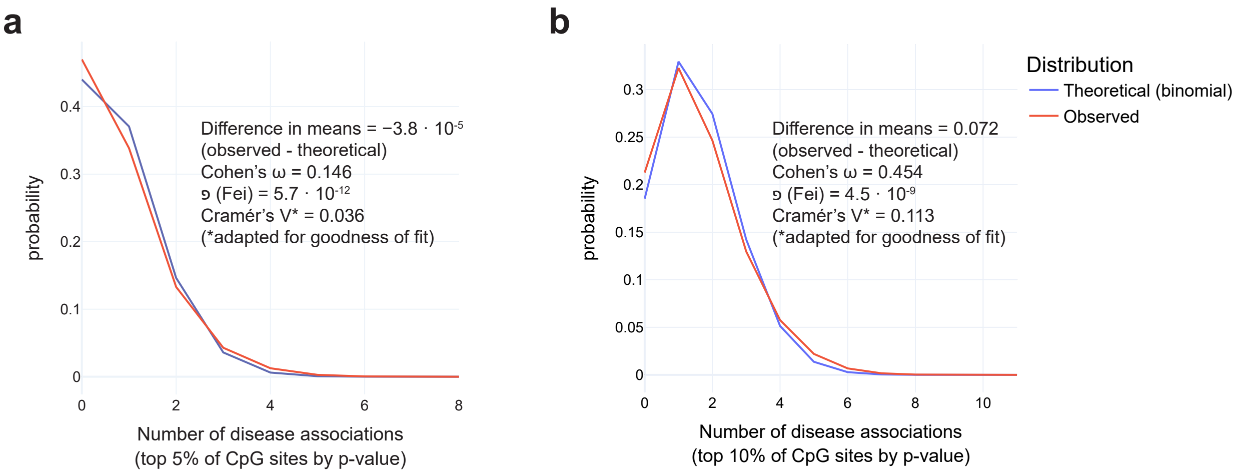


**Figure S8.** Distributions of the number of times a CpG site appeared in the top *N* % of CpG sites associated with human chronic diseases (16 diseases in total) across all CpG sites (more than 250,000 CpG sites). The observed distribution is compared to the theoretical binomial distribution with *n*=16 and *p*=0.01**N*. For details on the metrics we used for effect size estimation, see "Materials and Methods", section "Meta-analysis of human chronic diseases in the context of aging, early childhood development, sex differences and species longevity".

**a**, *N*=5.

**b**, *N*=10.

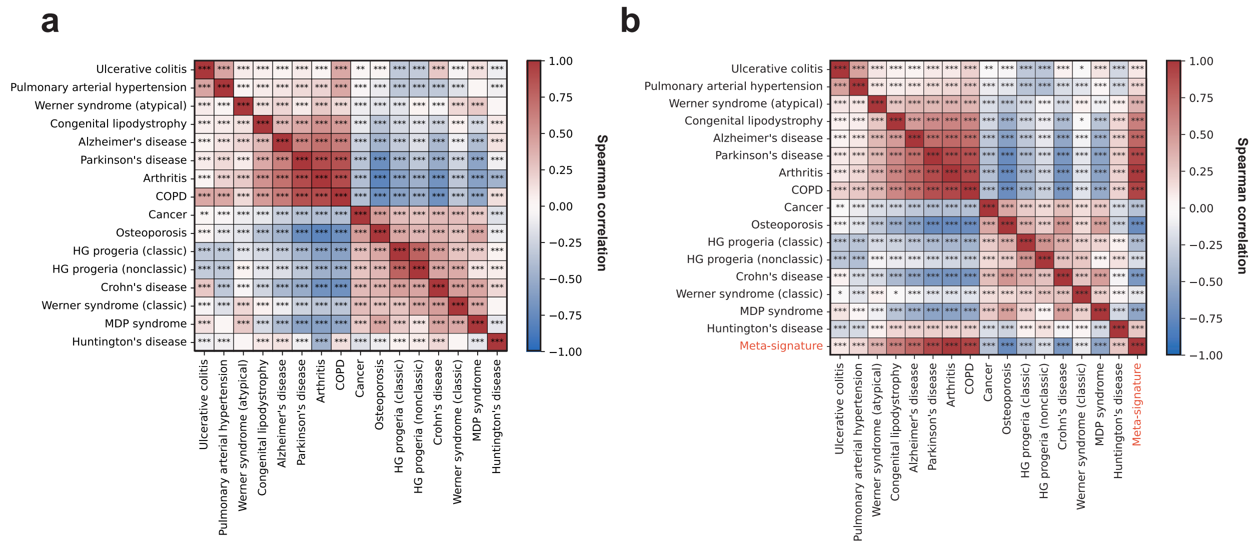


**Figure S9.** Spearman correlations of disease signatures calculated using intersections of top 100,000 CpG sites ranked by p-value (left column) and using top *N* CpG sites ranked by p-value in the meta-signature (right column), where *N* is equal to the mean intersection size from the corresponding plot from the left column.

**a**, Diseases only, only the CpG sites that were present in the meta-signature were used.

**b**, Diseases and their meta-signature, *N*=40,969.

ns – p adjusted ≥ 0.1 (not significant); ^ – p adjusted < 0.1, * – p adjusted < 0.05; ** – p adjusted < 0.01; *** – p adjusted < 0.001.

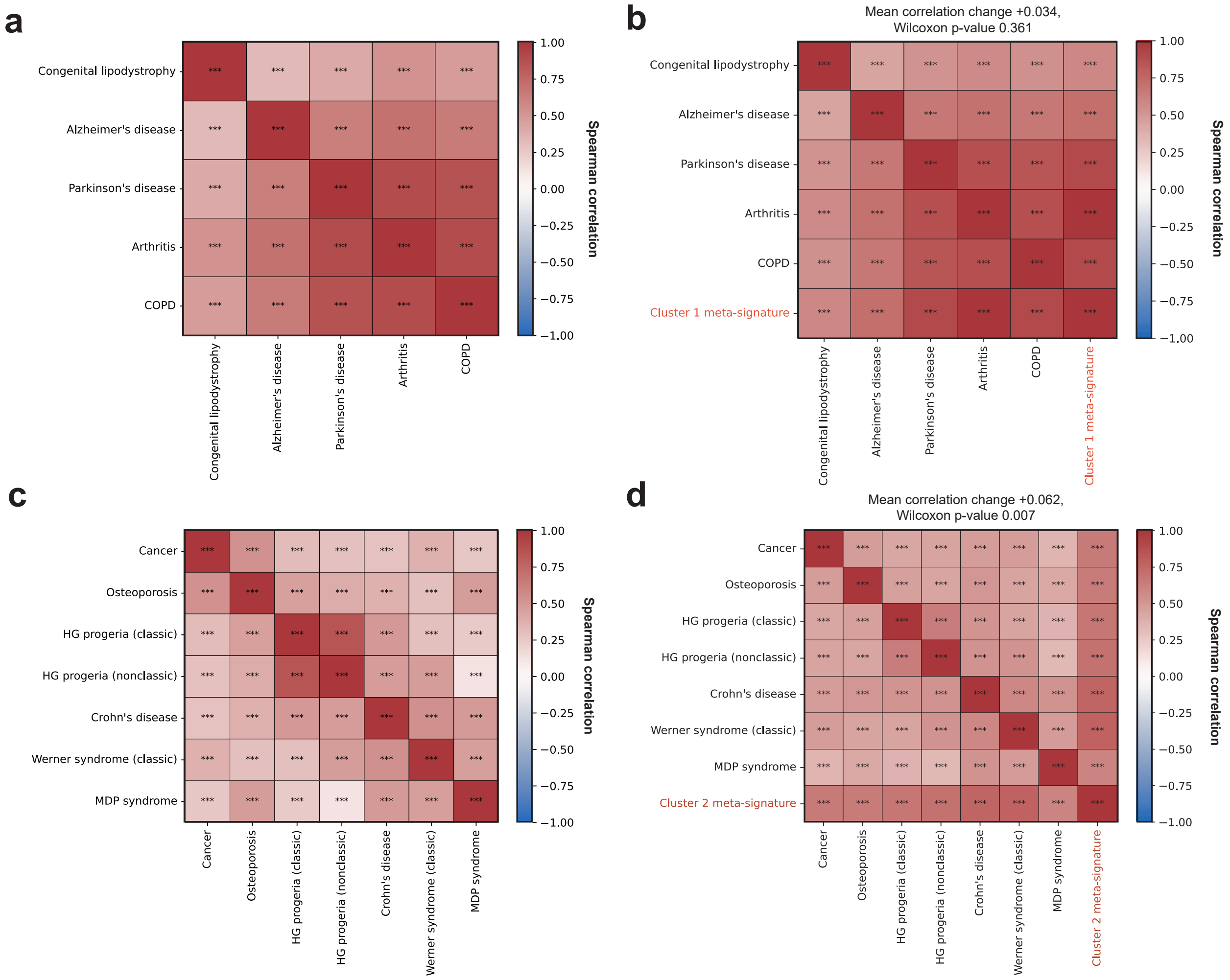


**Figure S10.** Spearman correlations of disease signatures calculated using intersections of top 100,000 CpG sites ranked by p-value in the corresponding disease signatures (left column) and using top *N* CpG sites ranked by p-value in the meta-signature (right column), where *N* is equal to the mean intersection size from the corresponding plot from the left column. In both columns, only the CpG sites that were present in the corresponding meta-signature were used.

**a**, Cluster 1 diseases.

**b**, Cluster 1 diseases and their meta-signature, *N*=52,166.

**c**, Cluster 2 diseases.

**d**, Cluster 2 diseases and their meta-signature, *N*=36,244.

ns – p adjusted ≥ 0.1 (not significant); ^ – p adjusted < 0.1, * – p adjusted < 0.05; ** – p adjusted < 0.01; *** – p adjusted < 0.001.

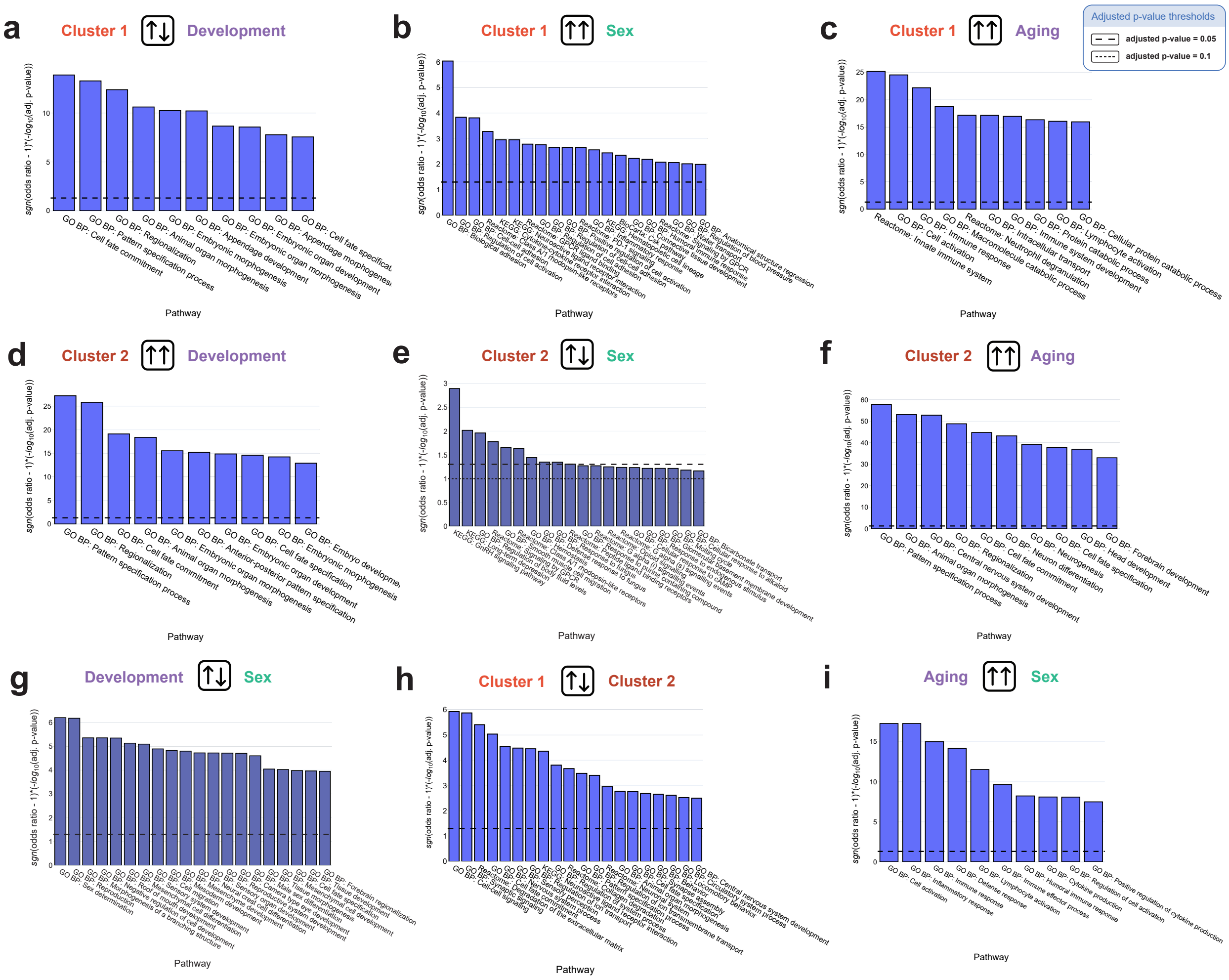


**Figure S11.** Intersections of CpG sites whose methylation changes in the same or opposite direction in two signatures with functional groups of genes. CpG sites were filtered by adjusted p-values in the respective signatures (we looked at the intersections of 100,000 CpG sites with lowest adjusted p-values). Intersection p-values were calculated using Fisher's exact test.

**a**, CpG sites whose methylation changed in the opposite direction in the meta-signature of cluster 1 diseases and the signature of early childhood development.

**b**, CpG sites whose methylation changed in the same direction in the meta-signature of cluster 1 diseases and the sex (male vs. female) signature.

**c**, CpG sites whose methylation changed in the same direction in the meta-signature of cluster 1 diseases and the signature of adulthood chronological age.

**d**, CpG sites whose methylation changed in the same direction in the meta-signature of cluster 2 diseases and the signature of early childhood development.

**e**, CpG sites whose methylation changed in the opposite direction in the meta-signature of cluster 2 diseases and the sex (male vs. female) signature.

**f**, CpG sites whose methylation changed in the same direction in the meta-signature of cluster 2 diseases and the signature of adulthood chronological age.

**g**, CpG sites whose methylation changed in the opposite direction in the signature of early childhood development and the sex (male vs. female) signature.

**h**, CpG sites whose methylation changed in the opposite direction in the meta-signature of cluster 1 diseases and the meta-signature of cluster 2 diseases.

**i**, CpG sites whose methylation changed in the same direction in the signature of adult chronological age and the sex (male vs. female) signature.

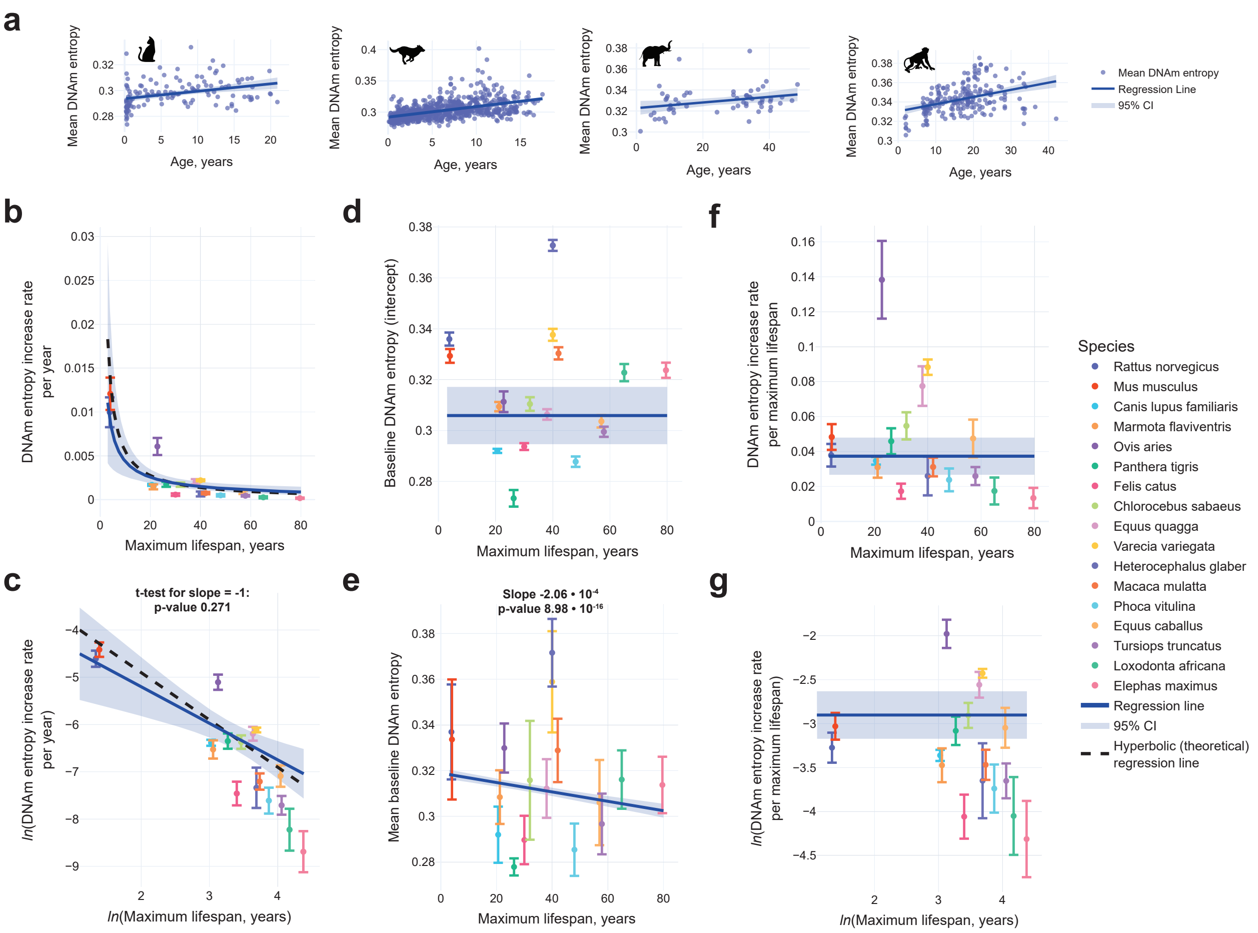


**Figure S12.** DNAm entropy associations with maximum lifespan on blood data. Error bars denote ±SD (standard deviation) in e, and ±SE (standard error) in all other subfigures. 95% CI stands for 95% confidence interval. The theoretical hyperbolic expected functional relationship between rate of change (ROC) per year and maximum lifespan is taken from Horvath et al.[1]. All regression lines were fitted with WLS (weighted least squares) with $\frac{1}{SE^2}$ weights, except for the regression line from e,, which was fitted with OLS (ordinary least squares) to raw mean DNAm entropy data.

**a**, Examples of regressions of mean DNAm entropy against chronological age (in years) for *Felis catus*, *Canis lupus familiaris*, *Loxodonta africana* and *Macaca mulatta* (left to right). Mean DNAm entropy is the same as total DNA methylation entropy defined in Materials and Methods. In this analysis, the entropy was computed without filtering CpG sites based on any association with maximum lifespan or chronological age.

**b**, DNAm entropy increase rate (slope) from per-species regressions of mean DNAm entropy against age (in years), plotted against maximum lifespan. The regression curve, 95% CI and theoretical regression curve are the transformed versions of the regression lines from c,.

**c**, Same as Fig. S12b, but in logarithmic coordinates. The p-value for the slope of the regression line is from a t-test with $H_0$: $slope = -1$ and $H_A$: $slope \neq -1$. For the theoretical regression line, the slope was fixed to -1 and only the intercept was fitted with WLS.

**d**, Intercept coefficients from per-species regressions of mean DNAm entropy against age (in years), plotted against maximum lifespan. WLS on intercepts did not result in a significant slope coefficient (p-value 0.51), so the regression line just represents the weighted mean (intercept-only WLS).

**e**, Mean baseline DNAm entropy against maximum lifespan. Here, the baseline DNAm entropy was calculated for each sample as $H_i - k_n * n_i$, where $H_i$ is the total (mean) DNAm entropy of the $i$-th sample, $n_i$ is its normalized age (age divided by the corresponding maximum lifespan) and $k_n$ is the coefficient estimate for the normalized age from $H \sim \text{maximum\_lifespan} + \text{normalized\_age}$ OLS regression that was fitted to all samples. Baseline DNAm entropy is essentially the entropy that "a sample would have if its age was 0" according to the aforementioned regression model. The plot shows the mean baseline DNAm entropy ±SD for each species, and the regression line corresponds to $intercept + k_m * \text{maximum\_lifespan}$, where intercept and $k_m$ (coefficient for maximum lifespan) are taken from the same regression described above. The t-test p-value on the plot is also from this OLS regression.

**f**, Same as Fig. S12b, but for each species total DNAm entropy is regressed against normalized age. WLS on slopes did not result in a significant slope coefficient (p-value 0.504), so the regression line just represents the weighted mean (intercept-only WLS).

**g**, Same as Fig. S12f, but logarithmic coordinates. WLS on slopes in these coordinates did not result in a significant slope coefficient (p-value 0.27), so the regression line just represents the weighted mean (intercept-only WLS).

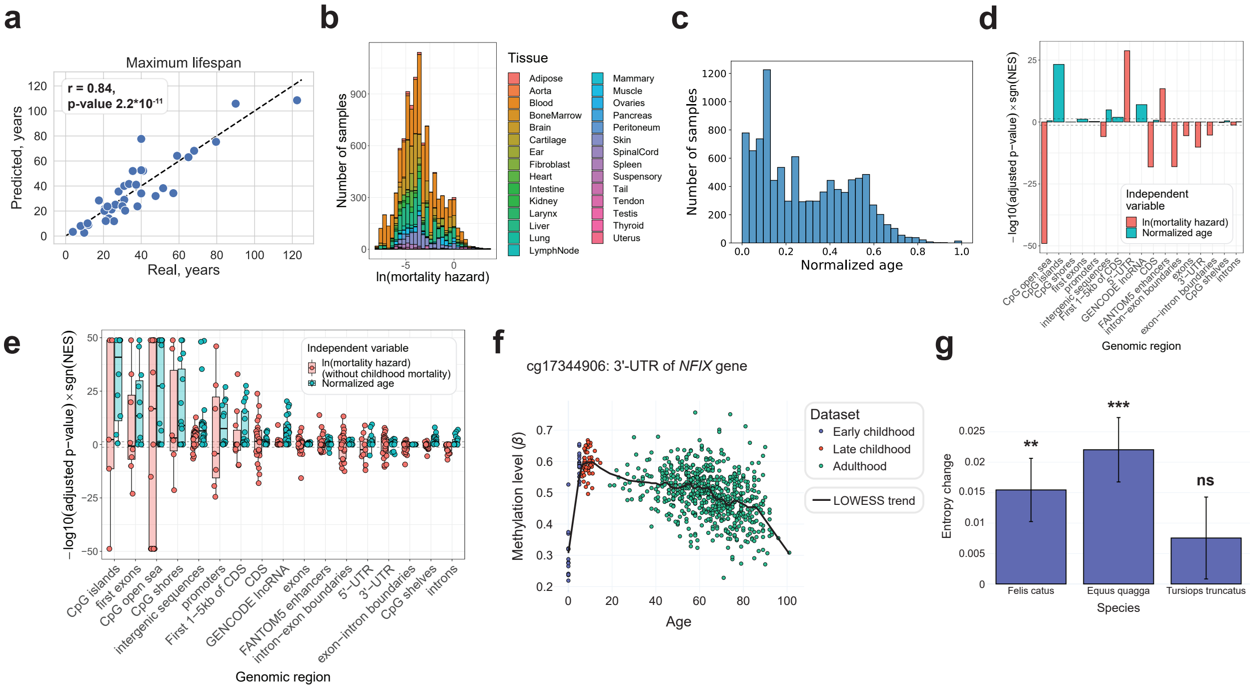


**Figure S13.** Mortality data and DNA methylation associations with mortality.

**a**, Real maximum lifespan against predicted, which was defined as the age at which survivorship function becomes less than 0.0001. Pearson correlation was 0.84, p-value $2.2 \cdot 10^{-11}$.

**b**, Distribution of mortality hazard by tissue.

**c**, Normalized age distribution.

**d**, GSEA enrichment of genomic regions in CpG sites associated with normalized age and mortality for all tissues and both sexes combined.

**e**, GSEA enrichment of genomic regions in CpG sites associated with normalized age and mortality, but mortality was calculated without the childhood mortality term. Dots represent combinations of tissue and sex for which the enrichment scores could be successfully calculated.

**f**, A second example of CpG sites whose methylation variation with age exhibits U-shaped behavior.

**g**, DNA methylation entropy changes during infant development (where the all-cause mortality declines) in the blood of 3 mammalian species: *Felis catus*, *Equus quagga* and *Tursiops truncatus*. The entropy was calculated for top 1000 CpG sites with the most significant methylation changes in infant development for each of the 3 species. ns – p-value ≥ 0.1 (not significant); ** – p-value< 0.01; *** – p-value < 0.001.

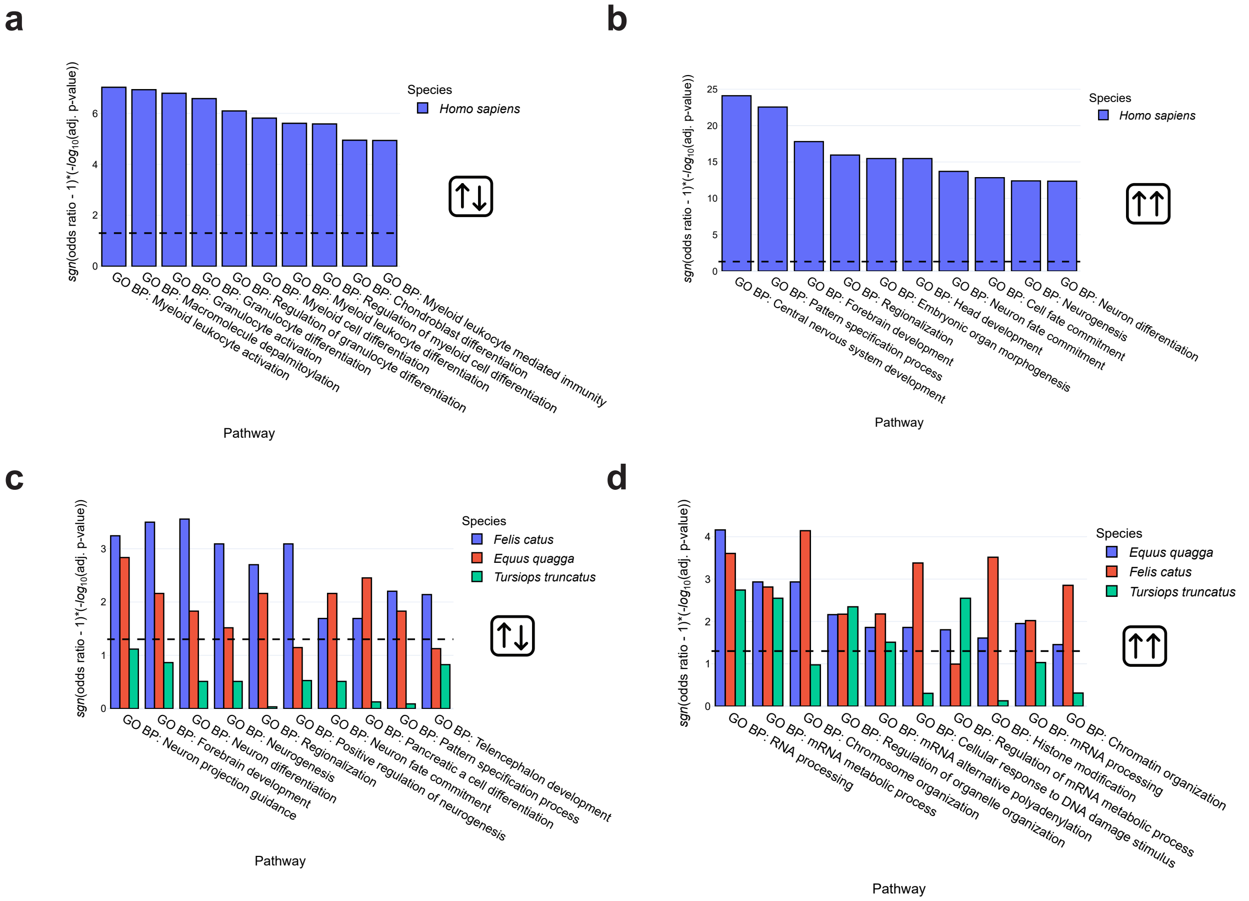


**Figure S14.** Fisher tests for intersections of functional groups of genes with CpG sites whose methylation variation with age exhibits monotonic and U-shaped behavior in blood data.

**a**, Results for human, CpG sites with U-shaped methylation changes.

**b**, Results for human, CpG sites with monotonically changing methylation.

**c**, Results for cat, plains zebra and common bottlenose dolphin, CpG sites with U-shaped methylation changes.

**d**, Results for cat, plains zebra and common bottlenose dolphin, CpG sites with monotonically changing methylation.

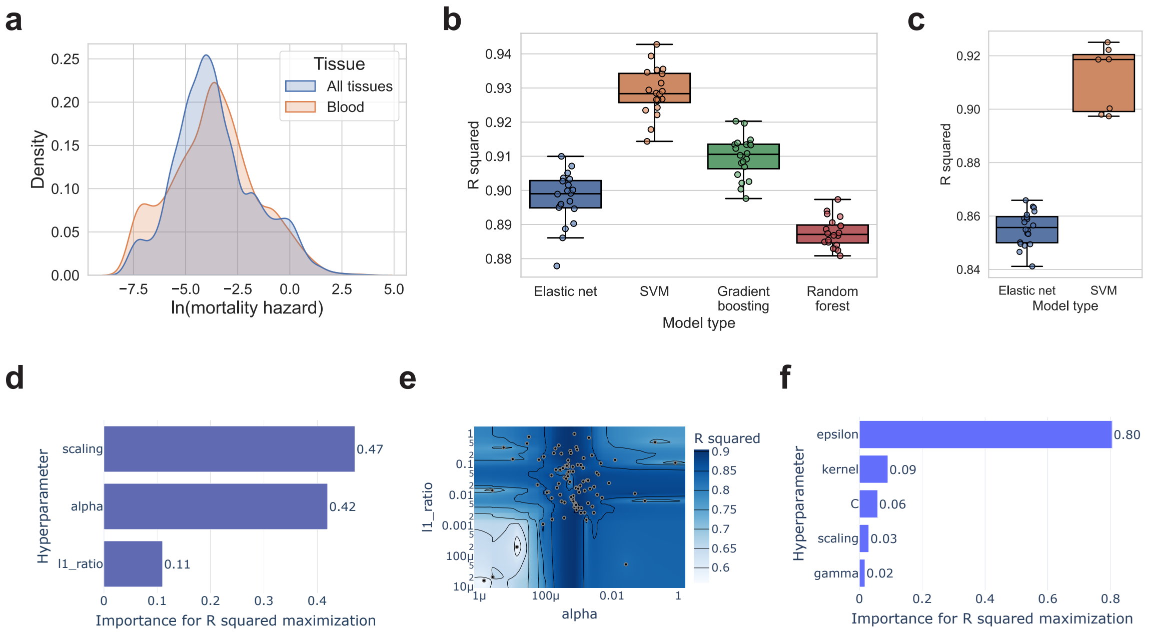


**Figure S15.** Target variable distributions and mortality clock model type comparison.

**a**, Target variable distributions for blood and all tissues.

**b**, Test set $R^2$ distributions for 4 model types for blood data.

**c**, Test set $R^2$ distributions for 2 model types for multiple tissues.

**d**, Hyperparameter importance for blood linear model clock from Bayesian hyperparameter optimization with optuna.

**e**, Contour plot for for blood linear model clock from Bayesian hyperparameter optimization with optuna.

**f**, Hyperparameter importance for blood SVM clock from Bayesian hyperparameter optimization with optuna.

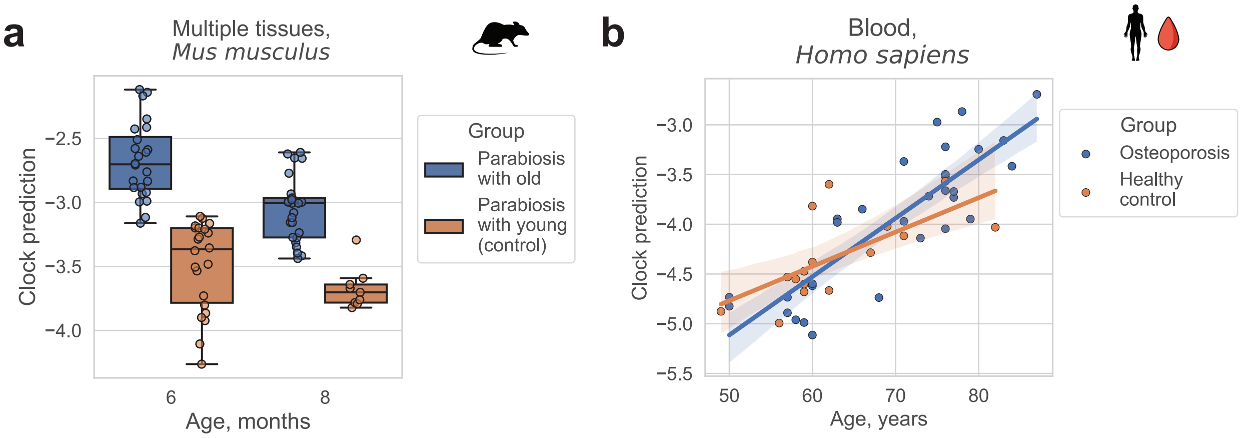


**Figure S16.** Examples of two types of disease or age-accelerating intervention effects.

**a**, Disease/intervention's effect on baseline predicted age is greater than the acceleration of predicted age increase with age. The clock is the multi-tissue elastic net linear model mortality clock. Mann-Whitney U test p-value for 6-month-old mice is $1.01 \cdot 10^{-8}$, for 8-month-old mice – $2.96 \cdot 10^{-5}$.

**b**, Disease/intervention's acceleration of predicted age increase with age is greater than the effect on baseline predicted age. The clock is the blood SVM mortality clock. OLS p-value of the group*age term coefficient is 0.06.

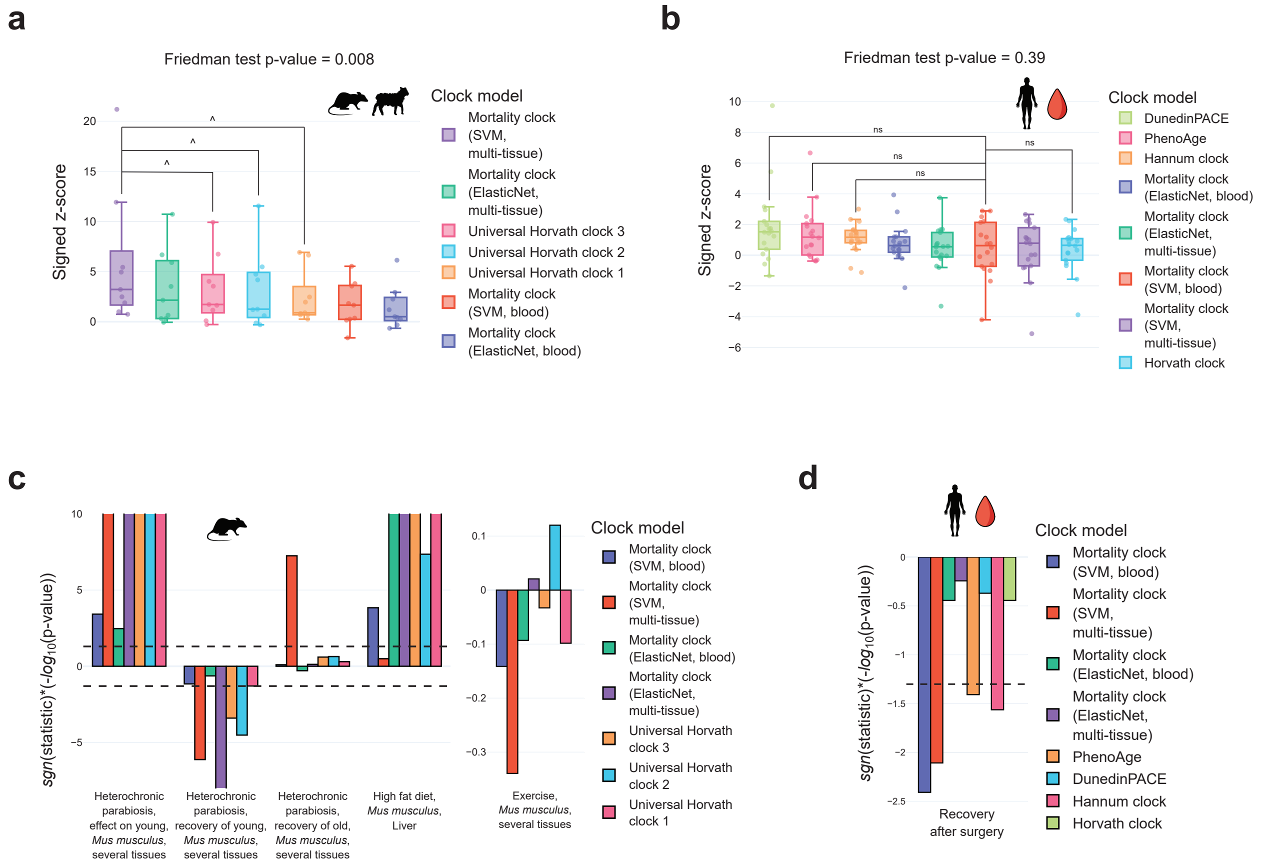


**Figure S17.** Mortality clock validation results on human blood data and on data from various tissues of model organisms (mice and sheep).

**a**, Mortality clocks vs. universal Horvath clocks on 9 mouse or sheep datasets from 6 studies. Statistical significance of the pairwise Wilcoxon tests is shown for the best mortality clock for blood data – the SVM model with RBF kernel. Friedman test was performed with all presented clocks as samples.

**b**, Mortality clocks vs. human benchmark clocks on 16 human chronic diseases and surgery. Statistical significance of the pairwise Wilcoxon tests is shown for the best mortality clock for multi-tissue data – the SVM model with RBF kernel. Friedman test was performed with all presented clocks as samples.

**c**, Mortality clocks vs. universal Horvath clocks on individual mouse datasets. The definition of “statistic” is equivalent to the one presented in Fig. 3b.

**d**, Mortality clocks vs. human benchmark clocks on recovery from hip trauma surgery. The definition of “statistic” is equivalent to the one presented in Fig. 3b.

ns – adjusted p-value ≥ 0.1, ^ – adjusted p-value < 0.1

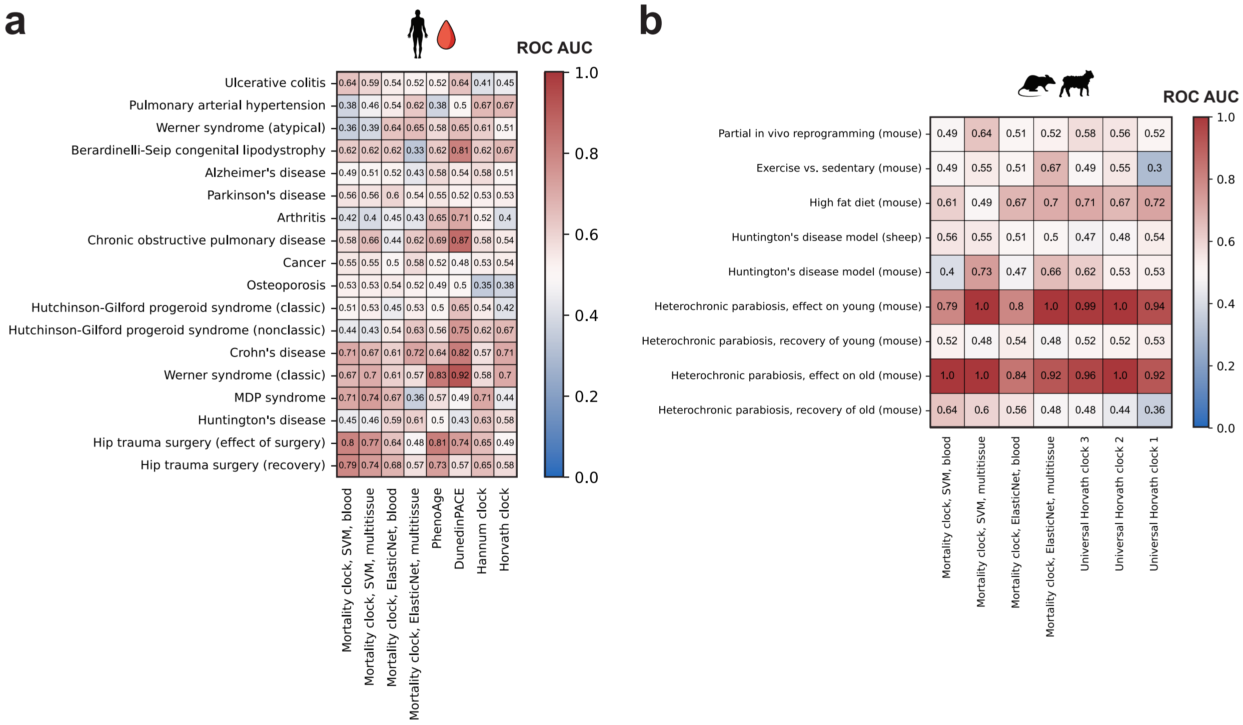


**Figure S18.** ROC AUC scores for age- and sex-adjusted epigenetic clock predictions for the classification of chronic disease patient samples and control group samples, as well as other conditions affecting biological age.

**a**, Human blood data.

**b**, Mouse and sheep data from blood and other tissues.
